# Paternal regulation of H3K4 methylation supports tumor suppressor networks in mammals intergenerationally

**DOI:** 10.64898/2026.08.28.747954

**Authors:** Benjamin William Walters, Rachel A. Heuer, Haoming Yu, Shubhangini Kataruka, Justin Tai, Katherine B. Henke, Zeyu Liu, Yadira M. Soto-Feliciano, Bluma J. Lesch

**Affiliations:** Department of Genetics, Yale School of Medicine, New Haven CT 06510; Keck Mass Spectrometry and Proteomics Resource, Yale School of Medicine, New Haven, CT 06510; Department of Biology, Massachusetts Institute of Technology, Cambridge MA 02139; Department of Obstetrics, Gynecology & Reproductive Sciences, Yale School of Medicine, New Haven CT 06510; Yale Cancer Center, Yale School of Medicine, New Haven CT 06510

## Abstract

Paternally-inherited epigenetic information can influence phenotype in offspring (1). Here, we identify a critical mechanistic contribution of KDM6A (UTX), an X-linked histone modifier and tumor suppressor, in regulating transmissible epigenetic information in mammalian sperm. Paternal loss of KDM6A increases cancer risk in genetically wild type offspring, but how *Kdm6a* knockout sperm transmit this effect at the molecular level is unknown (2). We find that KDM6A functions in spermatogenesis to promote methylation of histone H3 lysine 4 (H3K4) via selective interaction with the COMPASS complex methyltransferase KMT2C (MLL3). KMT2C and KDM6A are coordinately recruited to promoters of active genes in spermatogenic cells, contrasting with recruitment to intergenic enhancers in other cell types (3, 4). Loss of KDM6A disrupts H3K4 methylation at promoters of tumor suppressor genes in spermatogonia, and some of these defects persist in epididymal sperm and correspond to impaired expression in preimplantation embryos. These genes are also misregulated in normal and malignant hematopoietic tissue of genetically wild type offspring, indicating that impaired H3K4 methylation in KDM6A-deficient male germ cells may preferentially alter regulation of tumor suppressor gene networks in development across generations.

## Introduction

During mammalian spermatogenesis, paternal chromatin is extensively modified and reorganized, culminating in pervasive nucleosome eviction with retention of 1–15% of histones in spermatozoa (5). These retained histones are present at developmental promoters as well as distal intergenic regions (6–10), harbor post-translational modifications (5), and are detectable in zygotic chromatin after syngamy (11, 12), suggesting that they may contribute to a form of epigenetic memory that instructs gene regulation in the next generation beyond spermatogenesis (13, 14). On the other hand, paternal chromatin undergoes extensive reprogramming by maternal factors soon after fertilization to establish a new epigenetic landscape permissive for totipotency (15). The extent to which paternally-transmitted histones can bypass this reprogramming to influence the next generation remains an open question.

Parental exposure to environmental stressors can elicit phenotypes in the next generation in the absence of genetic alterations, implying that some paternally derived epigenetic memory can escape erasure. The best-studied mechanisms for this epigenetic transmission are small non-coding RNAs and DNA methylation in sperm. Several groups have independently shown that paternal metabolic stress can induce developmental abnormalities and adult-onset metabolic dysfunction in offspring, and have linked these transmitted phenotypes to alterations in sperm small RNAs (16–19). While less is known about the contribution of paternal histone modifications, some studies have implicated alterations in paternal H3K4 methylation in epigenetic transmission. For example, folate deficiency or overexpression of the H3K4 demethylase KDM1A in fathers yields sperm with altered H3K4me3 and is associated with skeletal deformities in the next generation (20, 21). However, more work is needed to understand how H3K4 methylation state is programmed in sperm and the mechanism by which alterations arise and elicit phenotypes in offspring.

H3K4 methylation is deposited by the multiprotein COMPASS (COMplex Proteins ASsociated with Set1) complex, which exists in multiple variants defined by mutually exclusive incorporation of one methyltransferase from among three paralogous pairs: SETD1A/B, KMT2A/B (MLL1/2), and KMT2C/D (MLL3/4). KMT2C/D-COMPASS includes KDM6A and NCOA6 as accessory subunits and is the primary variant for depositing H3K4me1 at gene-distal enhancers (22, 23), while variants defined by SETD1A/B and KMT2A/B associate with distinct accessory subunits, including CXXC1 and MENIN respectively, and primarily deposit H3K4me3 at promoters. Several COMPASS subunits, including KMT2B, the core subunit ASH2L, PTIP, and CXXC1 are essential for fertility in male mice largely due to their requirement in meiotic progression (24–27). The severity of these spermatogenic mutant phenotypes has precluded investigation of how their activities could affect chromatin in mature sperm and impact phenotype in the next generation.

We previously reported that deletion of the accessory COMPASS subunit KDM6A in the paternal germ line (*Kdm6a* cKO) leads to shortened lifespan and increased tumor burden in genetically wild type male offspring compared to genetically identical controls (2, 28–30). These findings suggest that loss of KDM6A alters epigenetic landscape in the paternal germ line and that some of these epigenetic changes are transmitted to the next generation, lowering the threshold for malignant transformation. Importantly, *Kdm6a* cKO males are fully fertile, enabling investigation of the mechanism by which epigenetic changes induced in sperm influence intergenerational phenotype (31). In addition to its non-catalytic function in H3K4 methylation as part of COMPASS, KDM6A is itself a histone modifier with catalytic demethylase activity for H3K27me3. However, we previously found only a modest effect on H3K27me3 in *Kdm6a* cKO male germ cells (2, 31), suggesting that the effects of KDM6A loss on offspring may be mediated primarily through its role in H3K4 methylation.

COMPASS components are among the most frequently mutated epigenetic regulators in human cancers. Deleterious mutations in *Kdm6a* are especially common in urothelial carcinoma, breast cancer, pancreatic cancer (32, 33), and leukemias (34), and numerous mechanistic studies have shown that dysregulation of KDM6A-containing COMPASS contributes to oncogenesis. Loss of *KDM6A* was reported to induce pancreatic cancer by deregulating COMPASS-mediated H3K4me1 and activating super-enhancers targeting oncogenes (35). KMT2C-KDM6A COMPASS also supports tumor suppressor gene expression programs in leukemic cells upon MENIN inhibition by enabling a molecular switch at the promoters of a common set of tumor suppressor genes (36). Furthermore, loss of KDM6A in hematopoietic stem cells altered gene expression, decreased H3K4me1, and affected ETS factor binding, ultimately shifting these cells towards a malignant state (3). A similar effect was observed in urothelial cancer, where loss of KMT2C/D reduced H3K4me1 and sensitized mouse urothelium to oncogenic transformation (37). Together, these studies point to functional loss of KDM6A-containing COMPASS as an early tumorigenic event that primes tissues for subsequent malignant transformation, reminiscent of the *Kdm6a* F1 phenotype.

Here, we sought an H3K27me3-independent pathway by which KDM6A loss in the male germline can confer cancer risk to male offspring. Surprisingly, we discovered that in male germ cells, KDM6A participates in COMPASS activity in a manner that differs qualitatively from somatic cells. In germ cells, KDM6A resides in the KMT2C-COMPASS variant and preferentially localizes to promoters of actively transcribed genes. Loss of KDM6A induced defects in H3K4 methylation beginning in spermatogonia that persisted into mature sperm. Genes retaining defective H3K4 methylation in sperm were enriched for cancer-related signaling pathways, and some of these were dysregulated in F1 embryos during preimplantation development and in malignant bone marrow cells of F1 animals. Many of these loci are also misregulated in *Kdm6a* mutant hematopoietic cells, revealing a KDM6A-sensitive tumor suppressor network responsive to intergenerational effects. Together, we find that KDM6A-containing KMT2C-COMPASS is an epigenetic regulator of heritable H3K4 methylation in the male germ line and highlights this pathway as a likely mechanism underpinning intergenerational inheritance of cancer risk.

## Results

### KDM6A interacts with KMT2C and the COMPASS complex in the male germ line

To discover how KDM6A regulates the epigenetic landscape during spermatogenesis, we performed an unbiased analysis of KDM6A protein partners in male germ cells by immunoprecipitating endogenous KDM6A from mouse testis tissue and performing mass spectrometry (IP-MS) to identify interacting proteins. Since *Kdm6a* transcript is expressed in a short developmental window during spermatogenesis, spanning late differentiating spermatogonia, meiotic entry, and early meiotic prophase (31), we first confirmed that KDM6A protein expression closely matches the *Kdm6a* transcript expression pattern, including in differentiating spermatogonia (KIT+), leptotene-zygotene spermatocytes (DMC1+) and Sertoli cells (Vimentin+) (**Fig. 1A, S1A**). Therefore IP-MS data from whole testis tissue is expected to capture interactors in late spermatogonia and early spermatocyte populations, as well as Sertoli cells.

**Figure 1.**
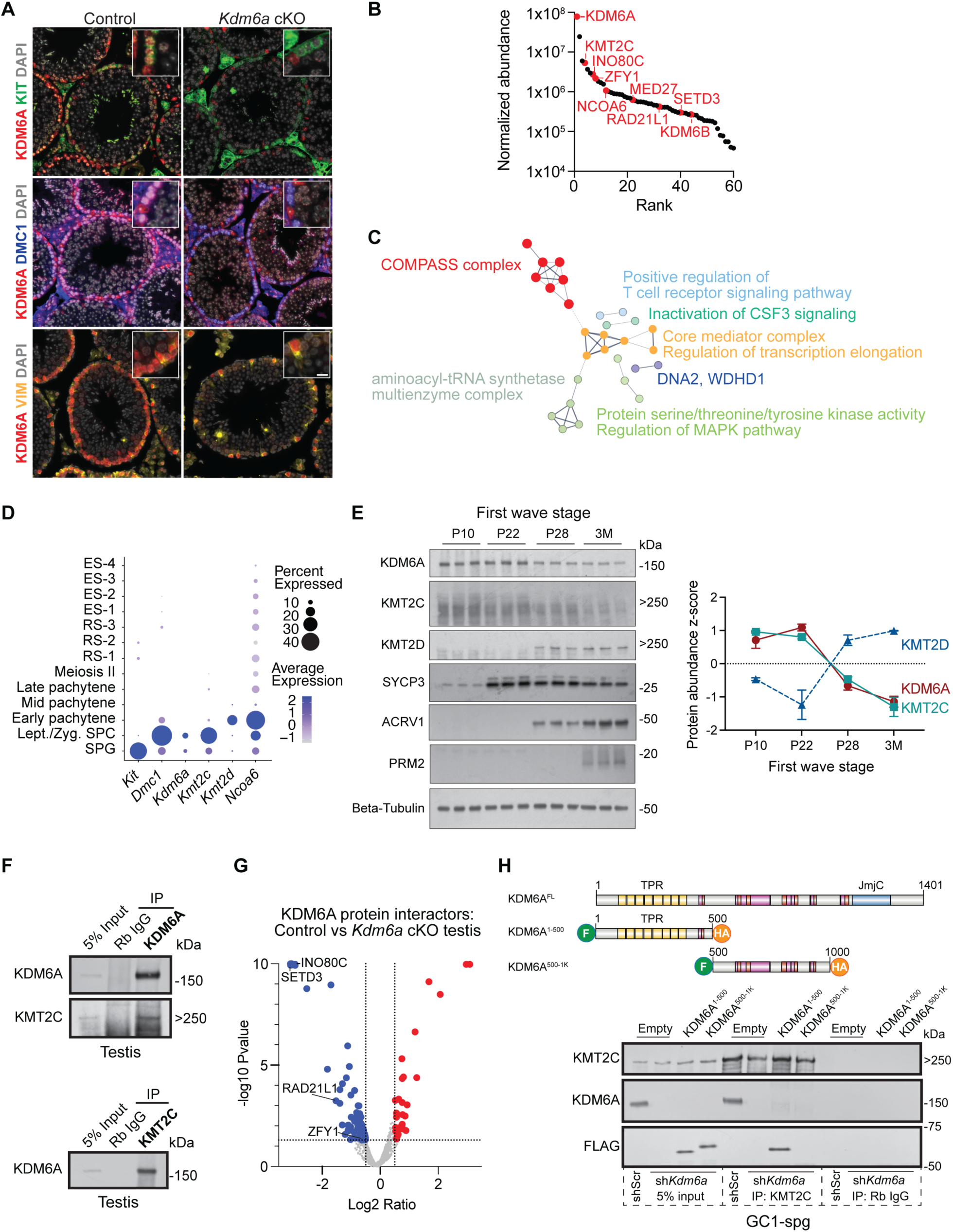
KDM6A interacts with COMPASS proteins in testis. **A,** Immunofluorescence for KDM6A and markers of differentiating spermatogonia (KIT), leptotene-zygotene spermatocytes (DMC1) and Sertoli cells (Vimentin) in control and *Kdm6a* cKO mouse testis sections. Scale bar = 10μm. **B,** Ranked normalized abundances for KDM6A protein interactors identified by mass spectrometry after immunoprecipitating endogenous KDM6A from whole mouse testis. Top interactors of KDM6A with potential functions in histone modification are highlighted in red. **C,** STRING network analysis for all KDM6A protein interactors identified in testis. Edges represent protein-protein interactions, and their thickness represents the strength of supporting data. Node colors and labels represent functional sets after K-means clustering. **D,** Gene expression for the indicated genes across spermatogenesis based on single cell RNA-seq in whole mouse testis (31). *Kit* is a marker for differentiating spermatogonia and *Dmc1* is a marker for leptotene/zygotene spermatocytes. SPG, spermatogonia; SPC, spermatocytes; RS, round spermatids; ES, elongating spermatids. **E,** Left, Western blot for KDM6A, KMT2C, and KMT2D in whole juvenile testes during the first wave of spermatogenesis representing addition of specific populations of spermatogenic cells: postnatal day 10 (P10, spermatogonia), P22 (spermatocytes, marked by SYCP3), P28 (elongating spermatids, marked by ACRV1), 3 months (spermatozoa, marked by PRM2). Right, quantitation of band intensity (z-score of log2 (target protein/tubulin). Error bars represent s.e.m. **F,** Western blot in whole mouse testes after immunoprecipitating KDM6A or KMT2C. Rb IgG, Rabbit IgG. **G,** Volcano plot showing differential KDM6A protein interactors between control and *Kdm6a* cKO mouse testes identified by label-free quantification mass spectrometry. Select factors discussed in the main text are indicated. **H,** Top, schematic of KDM6A protein domains and fragments expressed in the experiment. The domain architecture is based on UniProt annotation of the TPR domain (yellow), JmjC domain (blue), disordered regions (pink), and polar residues, low complexity regions, or basic and acidic residues (pink). Bottom, Western blot for FLAG, KMT2C, or KDM6A following immunoprecipitation with anti-KMT2C or Rabbit IgG in GC1-spg cells expressing shScr or sh*Kdm6a* along with one of two truncated fragments of KDM6A: 1-500 (amino acid 1-500) and 500-1K (amino acid 500-1000). Anti-KDM6A antibody recognizes the truncated C-terminus and therefore no band is detected in cells expressing only the KDM6A^1-500^ or KDM6A^500-1K^ fragments.

We qualitatively identified 121 candidate protein interactors for KDM6A using IgG as a negative control (see Methods, **Supplementary Table S1, Fig. S1B**). As expected, KDM6A itself was the most abundant protein recovered (**Fig. 1B**). COMPASS complex members overall were enriched among KDM6A interactors based on both STRING network analysis (38) and enrichment of Gene Ontology categories (39) (**Fig. 1C, S1C**). Members of the KMT2C-COMPASS complex variant (KMT2C and NCOA6) were among the top-ranked protein interactors, consistent with the interaction of KDM6A with this complex in other contexts (23, 40). Surprisingly, we did not identify the KMT2C paralog KMT2D as an interactor despite numerous reports of KDM6A also participating in KMT2D-defined COMPASS in other systems (41). This paralog specificity can be attributed to a developmental switch between paralogs, as both KMT2C mRNA and protein closely overlap the highly transient KDM6A expression pattern during spermatogenesis, while KMT2D is expressed later in development (**Fig. 1D-E, S1D-E**). Both KDM6A immunoprecipitation (IP) and reciprocal KMT2C IP in whole testis (**Fig. 1F**) or immortalized spermatogonia-derived GC1-spg cells (42) (**Fig. S1F**) confirmed the interaction between KDM6A and KMT2C (**Fig. 1F**).

In addition to COMPASS, KDM6A IP-MS identified several other chromatin-associated interactors, including INO80C, an ATP-dependent chromatin remodeler that has recently been reported to interact with COMPASS to mediate H3K4 methylation in *Arabidopsis* (43, 44), the KDM6A homolog KDM6B, and the putative histone methyltransferase SETD3, as well as the core mediator complex, DNA replication and damage response factors, and the aminoacyl tRNA synthetase complex (**Fig. 1B-C, S1C**). We also identified two testis-specific nuclear factors as KDM6A protein interactors: ZFY1, a Y-linked transcriptional activator (45); and RAD21L1, a meiosis-specific cohesin subunit (46). KDM6A therefore has both COMPASS and non-COMPASS chromatin functions in testis.

To identify KDM6A interactors specific to germ cells, we took advantage of our previously described *Kdm6a* conditional knockout (cKO) model where *Kdm6a* is deleted throughout postnatal spermatogenesis using a *Ddx4-Cre* transgene (2, 31). We collected KDM6A IP-MS data from *Kdm6a* cKO testes followed by label-free quantitative analysis in comparison to our wild type testis data. We verified substantially reduced KDM6A signal in *Kdm6a* cKO testis by Western blot (**Fig. S1B**) and based on normalized KDM6A abundance in the mass spectrometry data (**Fig. S1G**); we have previously shown that KDM6A protein is completely absent in isolated germ cells in the *Kdm6a* cKO model (31). As expected, more interactions were lost than gained in the *Kdm6a* cKO condition (n=111 lost vs. n=36 gained; **Fig. 1G, Supplementary Table S2**). INO80C and SETD3 showed the strongest reduction in the *Kdm6a* cKO condition and likely represent germ cell-specific KDM6A interactions (**Fig. 1G**). Appropriately, we also found loss of interaction with the germ cell-specific factors ZFY1 and RAD21L1 (**Fig. 1G**). We detected a trend toward reduced abundance of KMT2C in the *Kdm6a* cKO although this did not reach statistical significance, suggesting that KDM6A is likely a component of KMT2C-COMPASS in both germ cells and Sertoli cells in the testis (**Fig. S1H**).

Given the enrichment for COMPASS components, known participation of KDM6A in the COMPASS complex and previously reported roles for H3K4 methylation in epigenetic inheritance (14, 20, 47), we focused our attention on understanding KDM6A-containing COMPASS function in male germ cells. First, to define the KDM6A protein domain responsible for mediating interaction with COMPASS in the male germ line, we tested the interaction between KMT2C and truncated versions of KDM6A. Cancer-derived mutations in the N-terminal TPR domain of KDM6A have been shown to impair physical interactions with the KMT2C-COMPASS complex, highlighting this domain as crucial for KDM6A’s non-catalytic activity within COMPASS (48). We expressed truncated KDM6A constructs corresponding to the N-terminal TPR domain (amino acids 1–500) or the central region (500–1000) with FLAG/HA tags in GC1-spg cells with or without endogenous KDM6A (expressing either shScramble or sh*Kdm6a*). Both truncated isoforms and full-length KDM6A localized to the nucleus as expected (**Fig. S1I**). Immunoprecipitation of KMT2C followed by Western blotting confirmed its interaction with endogenous full-length KDM6A as well as with the KDM6A^1-500^ fragment, whereas no interaction was detected with the KDM6A^500-1K^ fragment (**Fig. 1H**). We conclude that KDM6A interacts with the COMPASS complex in male germ cells in part via interaction between the KDM6A N-terminal TPR domain and KMT2C.

### KDM6A is preferentially enriched at promoters in germ cells

To understand how KDM6A contributes to COMPASS function in the male germ line, we next evaluated the genomic distribution of KDM6A in spermatogenic cells and its effects on histone modification enrichment. We collected KDM6A ChIP-seq data in control and *Kdm6a* cKO testes (**Fig. S2A, S2B**). Over one quarter (n=12,680, 26%) of consensus KDM6A peaks were reduced or absent in *Kdm6a* cKO testes (**Fig. S2C**), suggesting that these peaks represent sites of KDM6A binding present predominantly or partially in germ cells (**Fig. 2A, S2D**). On the other hand, we infer that peaks unchanged in *Kdm6a* cKO testes represent KDM6A binding in somatic testicular populations, especially Sertoli cells where KDM6A is robustly expressed. To validate this assumption, we also collected KDM6A CUT&Tag data from isolated populations of differentiating spermatogonia (KIT+ cells; see Methods), which represent a subset of KDM6A+ germ cells. We confirmed high KDM6A CUT&Tag signal in the KDM6A ChIP-seq peaks that displayed differential enrichment between control and *Kdm6a* cKO, and virtually undetectable signal at ChIP-seq peaks that were unchanged in the *Kdm6a* cKO, consistent with these classes representing germ and somatic KDM6A binding, respectively (**Fig. 2B**).

**Figure 2.**
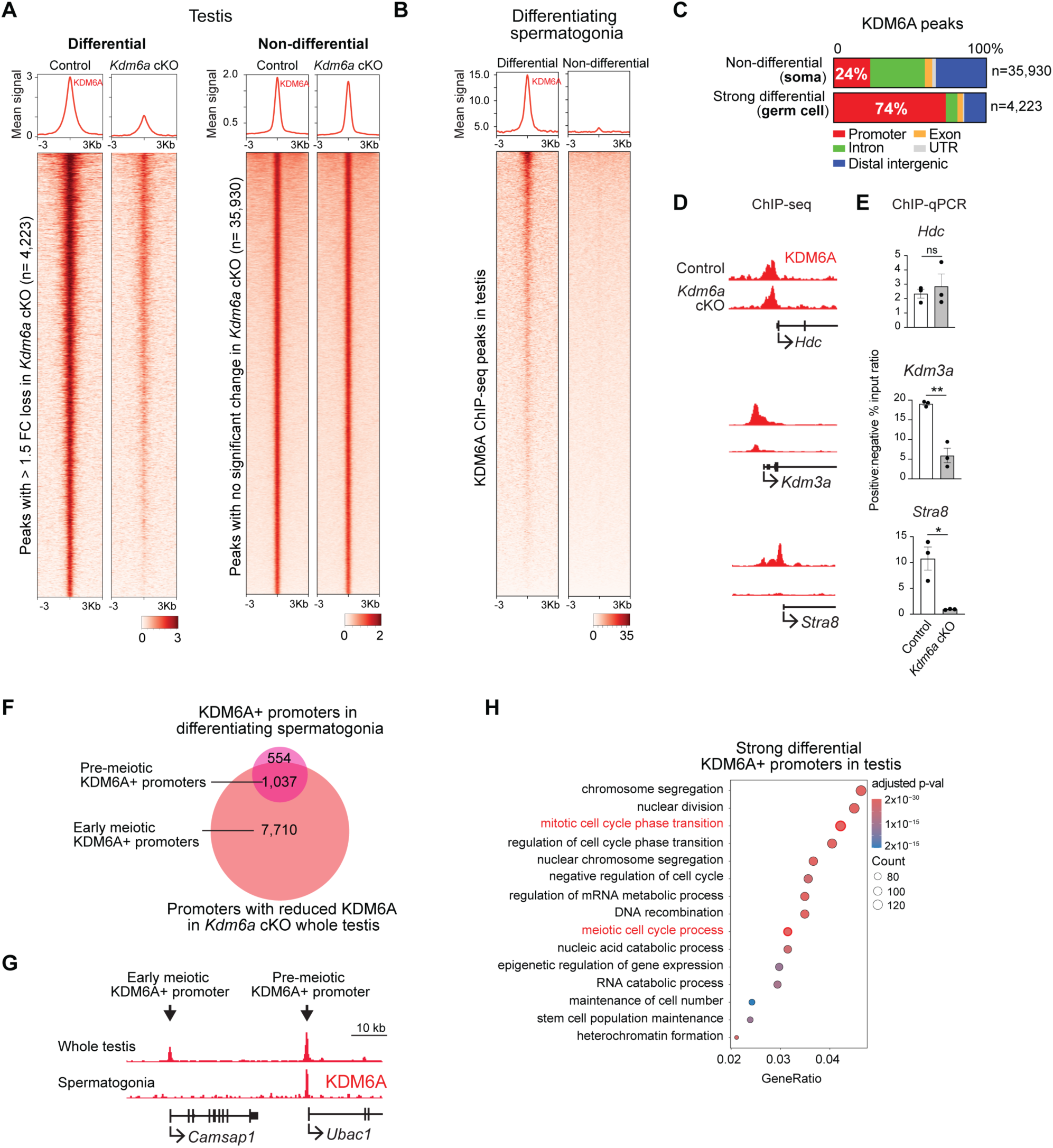
KDM6A predominantly binds promoters in male germ cells. **A.** RPKM-normalized signal for KDM6A ChIP-seq datasets in control and *Kdm6a* cKO mouse testes centered on peak summits. Left, peaks with >1.5 fold change difference in signal (FDR < 0.10) between control and *Kdm6a* cKO testes, inferred as germ cell binding events (n=4,223); right, peaks with no significant difference in signal between control and *Kdm6a* cKO testes, inferred as somatic binding events (n=35,930). **B,** RPKM-normalized signal for KDM6A CUT&Tag datasets in sorted KIT+ differentiating spermatogonia centered on KDM6A peaks identified as differential (germ cell) or non-differential (somatic) between control and *Kdm6a* cKO testis. **C,** Distribution of genome features for germ cell or somatic KDM6A peaks. **D,** Genome browser tracks for representative KDM6A promoter peaks from somatic (top), mixed germ cell/somatic (middle), and germ cell (bottom) classes in control and *Kdm6a* cKO whole testes. **E,** ChIP-qPCR for KDM6A at the promoters shown in (**D**). Points represent n=3 biological replicates and error bars show standard error of the mean. *p<0.05, **p<0.01, n.s. not significant, unpaired t-test. **F,** Overlap between gene promoters with inferred germ cell KDM6A ChIP-seq peaks in whole mouse testis (panel **A, Fig. S2D**) and promoters with KDM6A CUT&Tag peaks in sorted differentiating spermatogonia (**B**). **G,** Genome browser tracks for KDM6A ChIP-seq and CUT&Tag data showing a representative region in which both a pre-meiotic and an early meiotic KDM6A peak is present. **H,** Enriched Biological Process (BP) GO categories among inferred germ cell-specific KDM6A+ promoters from ChIP-seq data.

In most cells, KDM6A-containing KMT2C- and KMT2D-COMPASS complexes localize primarily to intergenic enhancer regions and regulate their activity by catalyzing monomethylation of lysine 4 on histone H3 (H3K4me1) and recruiting p300/CBP to promote acetylation of H3K27 (H3K27ac) (4, 35, 49–52). Unexpectedly, we observed a striking difference in genomic distribution between differential (germ cell) and non-differential (somatic) testicular KDM6A peaks. Whereas the majority of somatic peaks were found at intronic or intergenic regions as expected, the majority of germ cell KDM6A peaks (74%) were at gene promoters (**Fig. 2C, S2E**). This observation suggests that KDM6A unexpectedly occupies promoters rather than enhancers in male germ cells, contrasting with its predominant recruitment to intronic and intergenic sites in somatic cells (53, 54). We validated enrichment of KDM6A at representative promoters in all three classes (somatic, **Fig. 2A**; somatic + germ cell [<1.5FC in *Kdm6a* cKO], **Fig. S2D**; germ cell [>1.5FC in *Kdm6a* cKO], **Fig. 2A**) by ChIP-qPCR in control and *Kdm6a* testes (**Fig. 2D-E**).

KDM6A peaks identified by CUT&Tag in spermatogonia were also frequently at promoters, and the majority (65%) were classified as germ cell-specific KDM6A+ promoters in our ChIP data (**Fig. 2F**). Meanwhile, germ cell KDM6A promoter ChIP-seq peaks that did not have a peak in spermatogonia CUT&Tag data (**Fig. 2F-G**) likely represent KDM6A binding events in early meiotic cells, which are not captured by sorting using the KIT surface marker. Indeed, genes with germ cell-specific KDM6A binding in whole-testis ChIP-seq data were strongly enriched for gene ontology terms related to both mitotic and meiotic cell cycle processes, in keeping with KDM6A expression at the transition between mitosis and early meiotic prophase (**Fig. 2H**), while genes with KDM6A promoter peaks in spermatogonia CUT&Tag data were not enriched for meiotic cell cycle processes and instead were associated with mitotic cell cycle functions as well as RNA regulation and DNA replication and repair (**Fig. S2F**). All together, these findings support a promoter-specific function for KDM6A in male germ cells.

### KDM6A co-occupies active promoters with KMT2C in male germ cells

Given their interaction in germ cells, we next asked whether KDM6A and KMT2C occupy the same genomic regions, supporting a coordinated function. ChIP-seq for KMT2C in whole testis revealed a distribution of peaks closely resembling that of KDM6A, with most peaks residing at promoters (69%, **Fig. 3A**). There was a striking overlap between promoters bound by KDM6A in germ cells (KDM6A signal either fully or partly lost in *Kdm6a* cKO testis) and promoters bound by KMT2C, suggesting that KDM6A may function as part of the KMT2C-COMPASS variant at promoters (**Fig. 3B, S3A**). We designated KMT2C+ promoters with germ cell-specific KDM6A occupancy as high-confidence promoters likely to be regulated by KDM6A-containing KMT2C-COMPASS in male germ cells, and selected these for further analysis (n=2,601, **Fig. S3A**, **Supplementary Table S3**).

**Figure 3.**
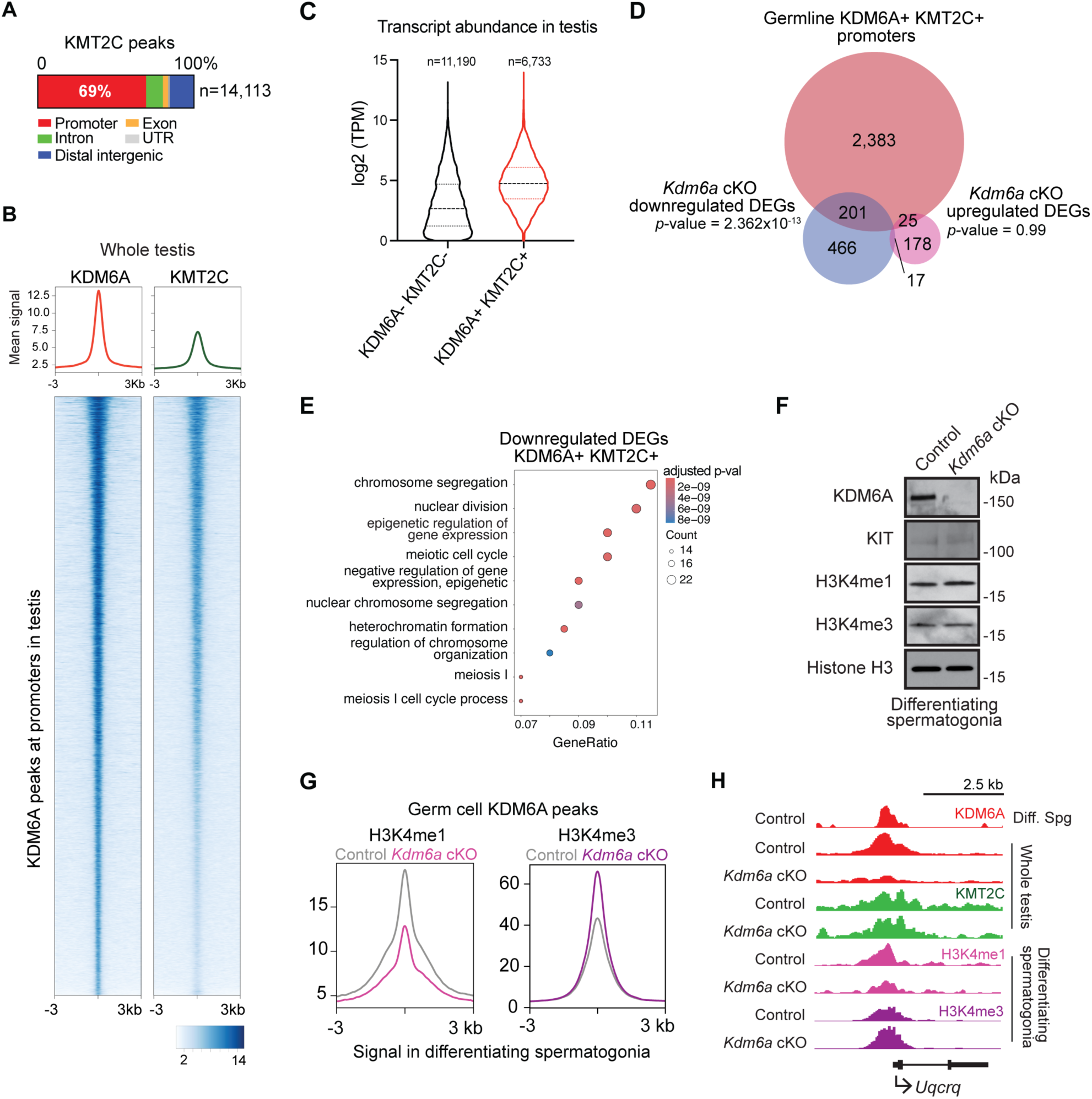
KDM6A coordinates with KMT2C to regulate H3K4 methylation at promoters. **A,** Distribution of genome features for KMT2C ChIP-seq peaks in mouse testis. **B,** RPKM-normalized signal for KDM6A and KMT2C ChIP-seq centered on KDM6A peaks at promoters. **C,** Transcripts per million (TPM) in whole mouse testis for genes with and without KDM6A promoter binding. **D,** Overlap of gene promoters co-occupied by KDM6A and KMT2C with differentially expressed genes identified by scRNA-seq in *Kdm6a* cKO testis (31). p-values calculated by Fisher’s Exact test. **E,** Enriched GO categories among downregulated genes in *Kdm6a* cKO testis whose promoters are bound by both KDM6A and KMT2C. **F,** Western blot for H3K4me1 and H3K4me3 in differentiating spermatogonia (KIT+) derived from control and *Kdm6a* cKO testes. Histone H3 is a loading control. **G,** RPKM-normalized signal for H3K4me1 and H3K4me3 in control and *Kdm6a* cKO differentiating spermatogonia centered on inferred germ cell KDM6A ChIP-seq peaks. **H,** Genome browser tracks at a representative promoter bound by KDM6A and KMT2C, and with reduced H3K4me1 and increased H3K4me3 signal in differentiating spermatogonia of *Kdm6a* cKO testes.

In other cell types, KDM6A is required to recruit the KMT2C paralog KMT2D to enhancers (4, 55). To assess if KDM6A plays an analogous role in recruiting KMT2C to promoters in male germ cells, we compared the binding profiles of KMT2C in *Kdm6a* cKO and control testes. We detected a slight but non-significant reduction in average KMT2C signal at genomic intervals bound by KDM6A in the male germ line, and a similar effect by ChIP-qPCR at a representative promoter (**Fig. S3B-C**). Therefore, while KDM6A co-localizes with KMT2C at promoters in male germ cells, it is largely dispensable for KMT2C recruitment.

Next, we assessed the relationship between KDM6A-KMT2C occupancy at promoters and transcriptional output of the associated genes. Genes whose promoters were bound by KDM6A-KMT2C were significantly more highly expressed than genes without KDM6A-KMT2C (**Fig. 3C, S3D**), indicating that KDM6A-KMT2C binds mostly active genes in testis. Then we asked if KDM6A-KMT2C occupancy at promoters regulates expression of the corresponding genes by comparing the set of KDM6A-KMT2C promoters to sets of differentially expressed genes we previously identified by single cell RNA-seq in *Kdm6a* cKO testes (DEGs) (31). We observed a statistically significant overlap between KDM6A-KMT2C bound promoters and DEGs (**Fig. 3D, Supplementary Table S4**), suggesting that KDM6A could directly regulate expression of these genes. Notably, we observed a greater overlap for downregulated DEGs (30%) compared to upregulated DEGs (12%), supporting a primary function for KDM6A-KMT2C in promoting gene expression as expected. This set of downregulated KDM6A-KMT2C bound DEGs was significantly enriched for genes encoding histone modifiers and factors involved in meiotic progression (**Fig. 3E**). Interestingly, the set of putative regulatory targets includes the *Kmt2c* gene (**Fig. S3E**) with *Kmt2c* transcript exhibiting significant depletion in *Kdm6a* cKO testes (**Fig. S3F**), suggesting that KDM6A and KMT2C may participate in an autoregulatory loop in germ cells.

### KDM6A-KMT2C regulate H3K4 methylation in differentiating spermatogonia

Next, we sought to define the molecular output of KDM6A-containing KMT2C-COMPASS activity in male germ cells. The KMT2C-COMPASS variant typically deposits mono-methylation of H3K4 (H3K4me1) at enhancers. Our finding that KDM6A-KMT2C is primarily found at promoters in germ cells raised the possibility that it may also regulate the promoter-associated mark H3K4me3, either directly or indirectly consequent to alteration of H3K4me1. We therefore assayed effects on multiple H3K4 methylation states, as well as the direct KDM6A target H3K27me3, in male germ cells lacking KDM6A.

In immortalized testis-derived cell lines (GC1-spg and GC2-spd (42, 56)), depletion of KDM6A using shRNAs reduced global levels of H3K4me1, H3K4me2, and H3K4me3, but had no effect on H3K27me3 (**Fig. S3G**). On the other hand, treatment with the demethylase inhibitor GSK-J4, which targets the catalytic activity of both KDM6A and its paralog KDM6B, resulted in increased H3K27me3 signal in GC1-spgs, GC2-spds, and ex vivo testis culture without affecting H3K4 methylation, revealing that KDM6A and KDM6B redundantly regulate H3K27me3 (**Fig. S3H-I**). This finding is consistent with our previous observation that single knockout of KDM6A has minimal impact on H3K27me3 in germ cells (31).

In whole testis (**Fig. S3J**) and sorted KIT⁺ differentiating spermatogonia (**Fig. 3F**), there was no change in global levels of either H3K4 methylation or H3K27me3 in *Kdm6a* cKO relative to control. This result is consistent with observations in mouse embryonic stem cells, where loss of KDM6A likewise does not affect global H3K4 methylation, but instead leads to localized reductions (27, 57–60). However, CUT&Tag for H3K4me1 and H3K4me3 in differentiating spermatogonia from control and *Kdm6a* cKO testis revealed a notable loss of H3K4me1 and gain of H3K4me3 at loci corresponding to KDM6A peaks in germ cells (**Fig. 3G-H**), suggesting that KDM6A-containing KMT2C-COMPASS regulates H3K4 methylation at these sites.

We conclude that in male germ cells, KDM6A preferentially occupies transcriptionally active promoters along with KMT2C and regulates H3K4 methylation balance, promoting H3K4me1 while restraining H3K4me3. This mechanism supports expression of genes important for chromatin regulation and meiotic division in the male germ line. However, germ cell KDM6A-KMT2C also occupies promoters of many genes where it does not impact expression, implying that it plays a different role in reinforcing the epigenetic landscape at these promoters.

### Altered H3K4 methylation persists in Kdm6a cKO sperm at promoters of signaling genes

We next probed if the H3K4 methylation changes induced in *Kdm6a* cKO spermatogonia were retained in sperm. Consistent with our findings in testis, Western blotting revealed no global changes in H3K4me1, H3K4me3 or H3K27me3 in caudal sperm of *Kdm6a* cKO mice (**Fig. S4A**). We also detected no significant differences in total levels of protamine 2 (PRM2) or histone H3, indicating that the histone-to-protamine transition is grossly unaffected and consistent with normal fertility of *Kdm6a* cKO males (31) (**Fig. S4A**).

We then assessed localized changes in H3K4me1 and H3K4me3 in caudal sperm by ChIP-seq. Control and *Kdm6a* cKO sperm datasets for H3K4me1 and H3K4me3, but not H3K27me3, separated along principal component 1 (PC1) in a principal component analysis, suggesting a genotype-dependent difference in sperm H3K4 methylation signal (**Fig. S4B**). At KDM6A-bound genomic intervals overall there was a slight reduction and broadening for both H3K4me1 and H3K4me3 signals (**Fig. 4A**). A peak-by-peak quantitative comparison of enrichment identified 1.44% of total H3K4me1 peaks (**Fig. 4B**) and 4.05% of total H3K4me3 peaks (**Fig. 4C**) as significantly altered (FDR < 0.10) in *Kdm6a* cKO sperm, with a strong bias towards gain of H3K4me1 and loss of H3K4me3. Notably, almost all peaks with loss of H3K4 methylation in sperm (either H3K4me1 or H3K4me3) occurred at promoters (**Fig. 4D**). In contrast, peaks with gain of either H3K4me1 or H3K4me3 typically occurred outside of promoters at intronic and intergenic regions. Average H3K4me1 signal was reduced at peaks with significant loss, but not gain, of H3K4me3 in sperm (**Fig. 4E**), suggesting an overall loss of H3K4 methylation at these promoter regions. On the other hand, there was little change in H3K4me3 signal at regions with significantly altered H3K4me1 (**Fig. 4E**). A total of 99 genes had significant changes in both H3K4me1 and H3K4me3 at their promoters in *Kdm6a* cKO sperm, representing a modest but statistically significant overlap (**Fig. 4F**). In total, we conclude that loss of KDM6A during spermatogenesis disrupts H3K4 methylation in sperm, especially at gene promoters normally enriched for H3K4me3.

**Figure 4.**
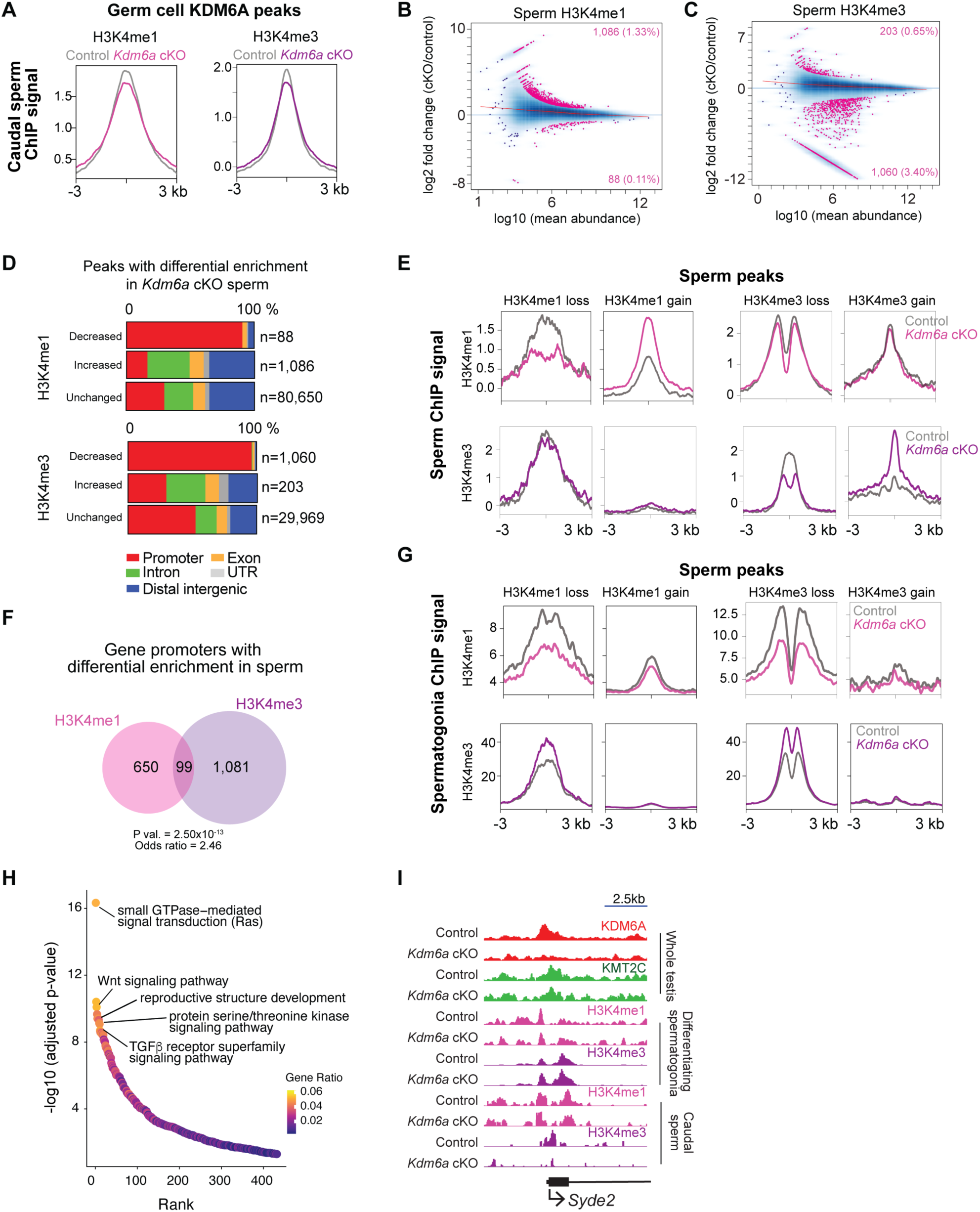
Sperm of *Kdm6a* cKO males retain perturbed H3K4 methylation at promoters. **A,** Mean H3K4me1 and H3K4me3 signal in control and *Kdm6a* cKO causal sperm centered on inferred germ cell KDM6A peaks from whole testis. **B-C,** Significant changes (pink, FDR < 0.10) in H3K4me1 (**B**) and H3K4me3 (**C**) signal in ChIP-seq data from *Kdm6a* cKO caudal sperm. **D,** Distribution of genome features for differential H3K4me1 and H3K4me3 peaks in *Kdm6a* cKO caudal sperm. **E,** Metagenes showing mean H3K4me1 (top) or H3K4me3 (bottom) signal in caudal sperm at peaks with differential H3K4me1 (left) or H3K4me3 (right) signal in *Kdm6a* cKO caudal sperm. **G,** Overlap between genes with differential H3K4me1 or H3K4me3 in *Kdm6a* cKO caudal sperm. P-value calculated by Fisher’s Exact test. **H,** Metagenes showing mean H3K4me1 (top) or H3K4me3 (bottom) signal in differentiating spermatogonia at peaks with differential H3K4me1 (left) or H3K4me3 (right) signal in *Kdm6a* cKO caudal sperm. **I,** Ranked plot showing enriched GO terms among genes whose promoters have significantly reduced H3K4me3 in *Kdm6a* cKO sperm. **J,** Genome browser tracks showing a representative promoter with reduced H3K4me1 and H3K4me3 in *Kdm6a* cKO sperm.

Since KDM6A is only transiently expressed during spermatogenesis in late spermatogonia and early meiotic cells (**Fig. 1A**), perturbation of H3K4 methylation in *Kdm6a* cKO sperm should be initiated by H3K4 methylation defects directly induced by KDM6A loss in these early cell types. We therefore compared the effects of *Kdm6a* cKO on H3K4me1 and H3K4me3 signal between sperm and differentiating spermatogonia (**Fig. 2B**). Average H3K4me1 signal was reduced in *Kdm6a* cKO spermatogonia at loci with any perturbation of H3K4me1 or H3K4me3 in *Kdm6a* cKO sperm, with the strongest effect at sites of either H3K4me1 or H3K4me3 loss in sperm (**Fig. 4G**). In contrast, average H3K4me3 signal was increased in *Kdm6a* cKO spermatogonia at loci with loss of either H3K4me1 or H3K4me3 in *Kdm6a* cKO sperm (**Fig. 4G**). These results imply that there is a relationship between KDM6A-dependent regulation of H3K4 methylation in spermatogonia and the level of H3K4me1 and H3K4me3 enrichment in sperm, although the direction of change in sperm may be affected by developmental events in the intervening period between KDM6A expression and sperm release.

Finally, we asked if perturbations in H3K4 methylation in sperm occur at genes that might have functional implications for development, tissue function, or tumorigenesis, given the cancer phenotype observed in offspring. We found that genes with significantly reduced promoter H3K4me3 in *Kdm6a* cKO sperm were strongly enriched for several developmentally important signaling pathways also implicated in cancer, including Wnt signaling, small GTPase signaling pathways including Ras, and TGF-beta signaling (**Fig. 4H-I**). In contrast, we did not observe any functional enrichment among genes with increased H3K4me3, or either increased or decreased H3K4me1 in *Kdm6a* cKO sperm. Interestingly, the promoters of the *Kmt2c* and *Kmt2d* genes both had reduced H3K4me3 in sperm, further suggesting a potential autoregulatory loop for the KDM6A-containing KMT2C-COMPASS complex. Notably, the widespread effects we observe on H3K4 methylation contrast with the more subtle effects on H3K27me3 signal we previously observed in *Kdm6a* cKO spermatogonia and sperm (2, 31). We conclude that alterations to H3K4 methylation induced by loss of KDM6A in the male germ line persist into mature sperm.

### Paternal loss of KDM6A alters gene expression in early preimplantation embryos

In order to affect gene regulation in adult somatic tissues, epigenetic information in *Kdm6a* cKO sperm must cross the barrier of epigenetic reprogramming that occurs soon after fertilization in preimplantation embryos. H3K4 methylation changes in sperm have been shown to impact gene regulation in developing embryos, suggesting that this information can in fact cross this barrier (14, 20). To directly test if germline loss of KDM6A can induce changes in gene regulation in the next generation, we assessed gene expression in embryos generated by *Kdm6a* cKO sperm. We collected RNA-seq data from pools of 2-cell and 8-cell embryos generated from control or *Kdm6a* cKO fathers naturally mated to wild type females. These two stages span a developmental window encompassing zygotic genome activation at the early 2-cell stage through establishment of the first major transcriptional program driving early embryonic cell fate specification. We identified 144 and 123 DEGs in 2-cell and 8-cell *Kdm6a* F1 embryos, respectively, with a 2-4-fold bias towards upregulation of expression at both stages (**Fig. 5A**). The size of the effect was greater at the 8-cell compared to the 2-cell stage, potentially reflecting amplification of early effects as development progresses. There was a small but statistically significant overlap between DEGs at the two stages (n = 18, **Fig. 5B**), indicating regulation of a common program.

**Figure 5.**
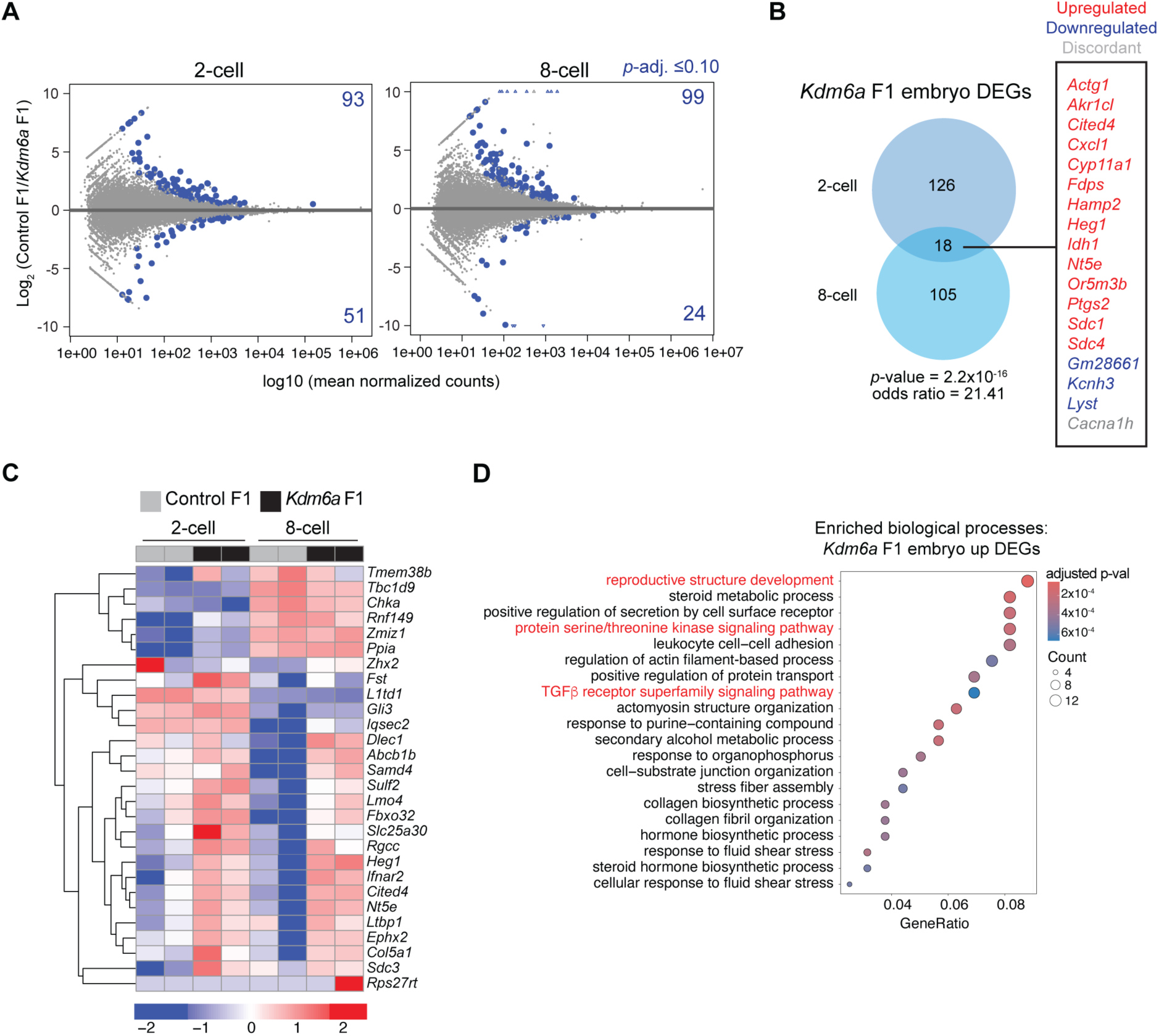
Altered expression in embryos derived from *Kdm6a* cKO sperm. **A,** MA plots for transcript levels in 2-cell and 8-cell embryos derived from *Kdm6a* cKO vs. control sperm. Differentially expressed genes (DEGs) are highlighted in blue. **B,** Overlap among DEGs at the 2-cell and 8-cell stages. **C,** Heatmap showing transcript level in embryos for genes called as DEGs in *Kdm6a* F1 embryos that also have differential histone methylation at their promoters in *Kdm6a* cKO sperm. **D,** Enriched Biological Process GO categories among transcripts upregulated in *Kdm6a* F1 embryos at either stage.

We asked whether paternally-derived alterations to H3K4 methylation could account for these changes in gene expression, and found that eleven percent of genes differentially expressed at either embryonic stage had significantly altered levels of H3K4 methylation in *Kdm6a* cKO sperm (n= 28/249, **Fig. 5C, S5A**). Notable genes in this set include tumor suppressors (*Zhx2* and *Dlec1*) and critical regulators of cancer development and progression (*Ltbp1* and *Lmo4*; **Fig. 5C**). Upregulated DEGs across both developmental stages were functionally enriched for “reproductive structure development” as well as the serine/threonine kinase signaling pathway and the TGFb pathway, pathways we also found to be enriched for differential H3K4 methylation in sperm (**Fig. 5D**). No functional enrichment was detected for downregulated DEGs.

Together, we conclude that regulatory changes are evident soon after fertilization in embryos generated by *Kdm6a* cKO fathers, and some of these correspond to genes where H3K4 methylation is disrupted in sperm. These findings indicate that some H3K4 methylation perturbations induced in *Kdm6a* cKO sperm may cross the fertilization barrier to impact gene regulation in offspring.

### KDM6A regulatory function in male germ cells parallels its tumor suppressive function in hematopoietic cells

Selective enrichment of KDM6A at promoters has been reported in leukemia cells (3, 36), reminiscent of our findings in male germ cells. In leukemia, KDM6A-containing COMPASS activity at promoters of tumor suppressor genes is important for their expression and antagonized by Menin, an alternative COMPASS subunit (36). If the tumor suppressor promoters regulated by KDM6A in leukemia are also KDM6A targets in germ cells, retained changes in H3K4 methylation in *Kdm6a* cKO sperm might preferentially impact regulation of these networks, increasing cancer risk. To further probe the similarities between KDM6A activity in germ and leukemic cells, we compared genome-wide occupancy of KDM6A in testis with KDM6A ChIP-seq datasets from mouse hematopoietic stem and progenitor cells (HSPCs) and from immortalized mouse leukemia cells (3, 36). Remarkably, we found extensive similarities in KDM6A binding between these cell types. Most KDM6A peaks were found at promoters in both HSPC and leukemia datasets (87% and 75%, respectively), similar to germ cells. Further, there was significant overlap between KDM6A+ promoters in HPSCs, leukemia cells, and male germ cells (n=3,934 germ cell KDM6A+ promoters, **Fig. 6A**). This set of shared target genes was strongly enriched for biological processes implicated in tumorigenesis such as cell division, RNA processing, and DNA repair as well as core and catalytic components of the COMPASS complex, including *Kmt2c* and *Kdm6a* itself (**Fig. S6A**). Motif analysis for each KDM6A ChIP-seq dataset, including our KDM6A CUT&Tag in spermatogonia, revealed a common and highly statistically significant enrichment for DNA binding motifs corresponding to the NFY, ETS and SP/KLF family of transcription factors (**Fig. 6B, S6B**). Notably, NF-YA and ELK4 have both been shown to cooperate with KDM6A to promote anti-leukemic gene expression programs (3, 36). Further, we previously detected a significant enrichment for ETS motifs at regions where differential DNA methylation persists from the *Kdm6a* cKO paternal germ line to *Kdm6a* F1 bone marrow cells (2). These findings reveal that a core set of target promoters important for tumor suppression is bound by KDM6A in both germ cells and hematopoietic cells.

**Figure 6.**
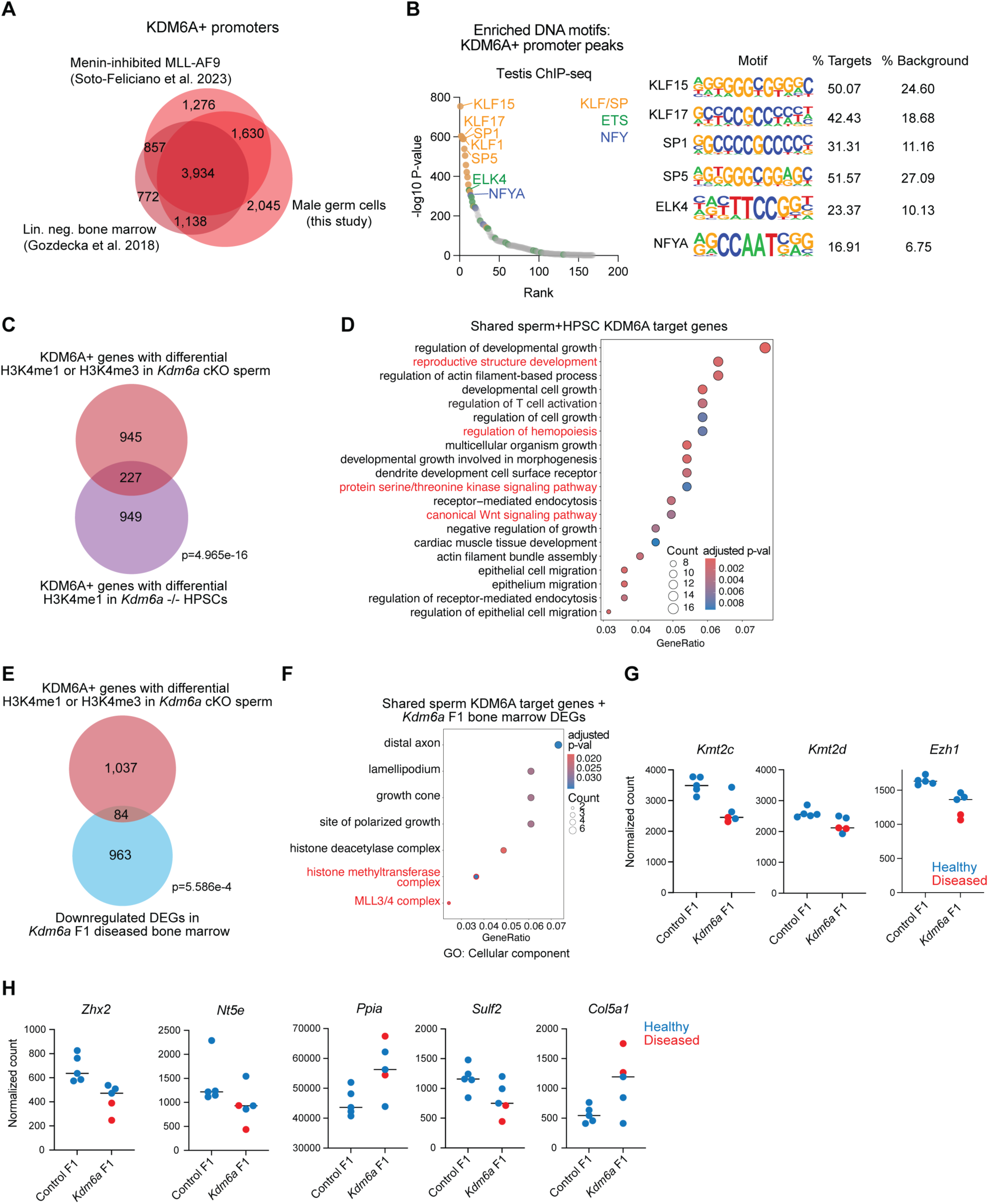
Shared regulatory activity of KDM6A in germ, hematopoietic, and leukemic cells. **A,** Overlap in gene promoters with KDM6A peaks in the male germline (this study), in HPSCs (3), and in leukemic cells treated with Menin inhibitor (36). **B,** Left, ranked plot showing enriched DNA motifs in KDM6A promoter-associated peaks in testis. Right, percent of germ cell promoter KDM6A peaks vs. background sequence containing top-ranked motifs. **C,** Overlap between genes with promoters bound by KDM6A and with differential H3K4 methylation in *Kdm6a* cKO sperm and genes with promoters bound by KDM6A with differential H3K4me1 in HPSCs. p-value calculated by Fisher’s Exact test. **D,** Enriched gene ontology categories among the 227 overlapping genes in (**C**). Terms highlighted in this study are in red. **E,** Overlap between genes with promoters bound by KDM6A and with differential H3K4 methylation in *Kdm6a* cKO sperm and downregulated genes detected in *Kdm6a* F1 malignant bone marrow. p-value calculated by Fisher’s Exact test. **F,** Enriched Cellular Component gene ontology categories among the 84 overlapping genes in (**E**). **G,** Library-normalized RNA-seq read counts for COMPASS-related genes in control F1 and *Kdm6a* F1 bone marrow. Data from histologically normal bone marrow is in blue; data from malignant bone marrow in red. **H,** Library-normalized RNA-seq read counts in control F1 and *Kdm6a* F1 bone marrow for genes differentially expressed in *Kdm6a* F1 embryos. Data from histologically normal bone marrow is in blue; data from malignant bone marrow in red.

To assess if H3K4 methylation at this shared set of KDM6A targets is sensitive to loss of KDM6A in both spermatogenic and hematopoietic cells, we next asked if the same alterations to H3K4 methylation detected in the *Kdm6a* cKO male germ line are also apparent in *Kdm6a* KO HSPCs. We focused our analysis on promoters bound by KDM6A in the male germ line and displaying differential H3K4 methylation in sperm, as these may represent sensitive loci directly regulated by KDM6A. We found a substantial and significant overlap between this set of genes and the set of promoters that is both bound by KDM6A in HSPCs and where H3K4me1 is perturbed in *Kdm6a* ^-/-^ HSPCs (**Fig. 6C**). This overlapping gene set represented 21% of genes with differential H3K4 methylation in *Kdm6a* cKO sperm (n=227 genes) and was enriched for functions relevant to tumorigenesis including the canonical Wnt signaling pathway, regulation of hematopoiesis, and serine-threonine kinase signaling pathway (**Fig. 6D**). Notably, this co-regulated gene set again includes *Kmt2c*.

Together, these findings indicate that altered H3K4 methylation in *Kdm6a* cKO sperm preferentially occurs at loci that are sensitive to KDM6A-dependent H3K4 methylation and linked to leukemogenesis in hematopoietic cells. Deleterious mutations in *Kdm6a* have been identified in mouse and human leukemias (34, 61–64), and deletion of *Kdm6a* in mouse HSPCs promotes a pre-leukemic state characterized by altered gene expression, localized gains in H3K27ac, and losses of H3K4me1, but notably no change in H3K27me3. These changes are consistent with a non-catalytic role for KDM6A in leukemogenesis (3, 65), and reminiscent of the function we identified for KDM6A and COMPASS in the male germ line. Interestingly, we previously observed a high incidence of blood cancers in *Kdm6a* F1 mice (2), raising the possibility that perturbation of shared KDM6A targets in sperm could establish a pre-sensitized epigenetic state in HSPCs analogous to direct mutation of *Kdm6a* itself.

### Genes sensitive to germline KDM6A-containing COMPASS activity have altered expression in blood tumors of offspring of *Kdm6a* cKO fathers

We next sought to determine if sperm-borne H3K4 methylation defects induced by loss of KDM6A could be linked to misregulation of these target genes in *Kdm6a* F1 offspring and contribute to tumorigenesis. We found a modest but statistically significant overlap (n=84) between KDM6A-bound genes with altered H3K4 methylation in *Kdm6a* cKO sperm and genes differentially expressed in bone marrow of *Kdm6a* F1 males harboring a myeloid-derived hematopoietic tumor, histiocytic sarcoma (2). This overlap was mostly driven by loci with reduced H3K4 methylation in *Kdm6a* cKO sperm and genes with reduced expression in *Kdm6a* cKO bone marrow (**Fig. 6E, S6C**). Interestingly, the set of 84 genes with reduced H3K4 methylation in *Kdm6a* cKO sperm and reduced expression in diseased *Kdm6a* F1 bone marrow was significantly enriched for H3K4 and H3K27 methyltransferase complex components, including *Kmt2c*, *Kmt2d* and *Ezh1* (**Fig. 6F, S6D**). Furthermore, although these genes were not differentially expressed at a statistically significant level in healthy *Kdm6a* F1 bone marrow, healthy *Kdm6a* F1 animals often exhibited reduced transcript levels for these genes, approaching statistical significance as a group (**Fig. 6G**).

We then examined whether genes differentially expressed in *Kdm6a* F1 embryos and associated with altered histone methylation in *Kdm6a* cKO sperm remain dysregulated in *Kdm6a* F1 bone marrow. Five genes from this set exhibited altered gene expression in *Kdm6a* F1 bone marrow harboring a tumor, including the tumor suppressor *Zhx2* (**Fig. 6H**). Again, although the expression of these genes was not statistically significantly altered as a group in healthy *Kdm6a* F1 bone marrow, some individuals exhibited large expression differences comparable to those observed in diseased samples.

Together, these results support a model where tumor suppressor genes normally regulated by KDM6A in both spermatogenic and hematopoietic cells inherit an abnormal epigenetic state at fertilization, mimicking depletion of KDM6A during hematopoiesis and lowering the threshold to transcriptional dysregulation and malignant transformation later in life. These dysregulated genes include *Kmt2c* and *Kmt2d*, suggesting that KDM6A coordinates a feedback loop reinforcing a stable H3K4 methylation landscape in both germ and hematopoietic cells and that faulty control of this regulatory machinery across generations may further exacerbate alterations in the histone methylation landscape to promote cancer.

## Discussion

Here, we identified a mechanism by which chromatin regulation in developing spermatogenic cells modulates intergenerational regulation of gene expression. KDM6A is recruited to promoters along with KMT2C, where it promotes H3K4 methylation. Loss of KDM6A results in alterations to H3K4me1 and H3K4me3 in differentiating spermatogonia that persist into mature sperm and preferentially occur at promoters of tumor suppressors and genes in signaling pathways implicated in cancer. Genes with defective H3K4 methylation in sperm exhibit dysregulated expression in preimplantation embryos and in healthy and diseased bone marrow of adult *Kdm6a* F1 offspring. KDM6A regulates highly concordant sets of genes in male germ cells, in hematopoietic cells, and in leukemia, making these shared targets especially vulnerable to KDM6A loss in the paternal germ line. These results nominate dysregulation of H3K4 methylation mediated by KDM6A-KMT2C COMPASS as a candidate mechanism for increasing cancer risk in genetically intact offspring of *Kdm6a* cKO males. We posit that alterations to H3K4 methylation in *Kdm6a* cKO sperm initiate a subtly altered epigenetic state at KDM6A-sensitive promoters including tumor suppressors in the next generation, conferring a suboptimum regulatory state at these genes in hematopoietic cells that lowers the threshold for tumorigenesis (**Fig. 7**).

**Figure 7.**
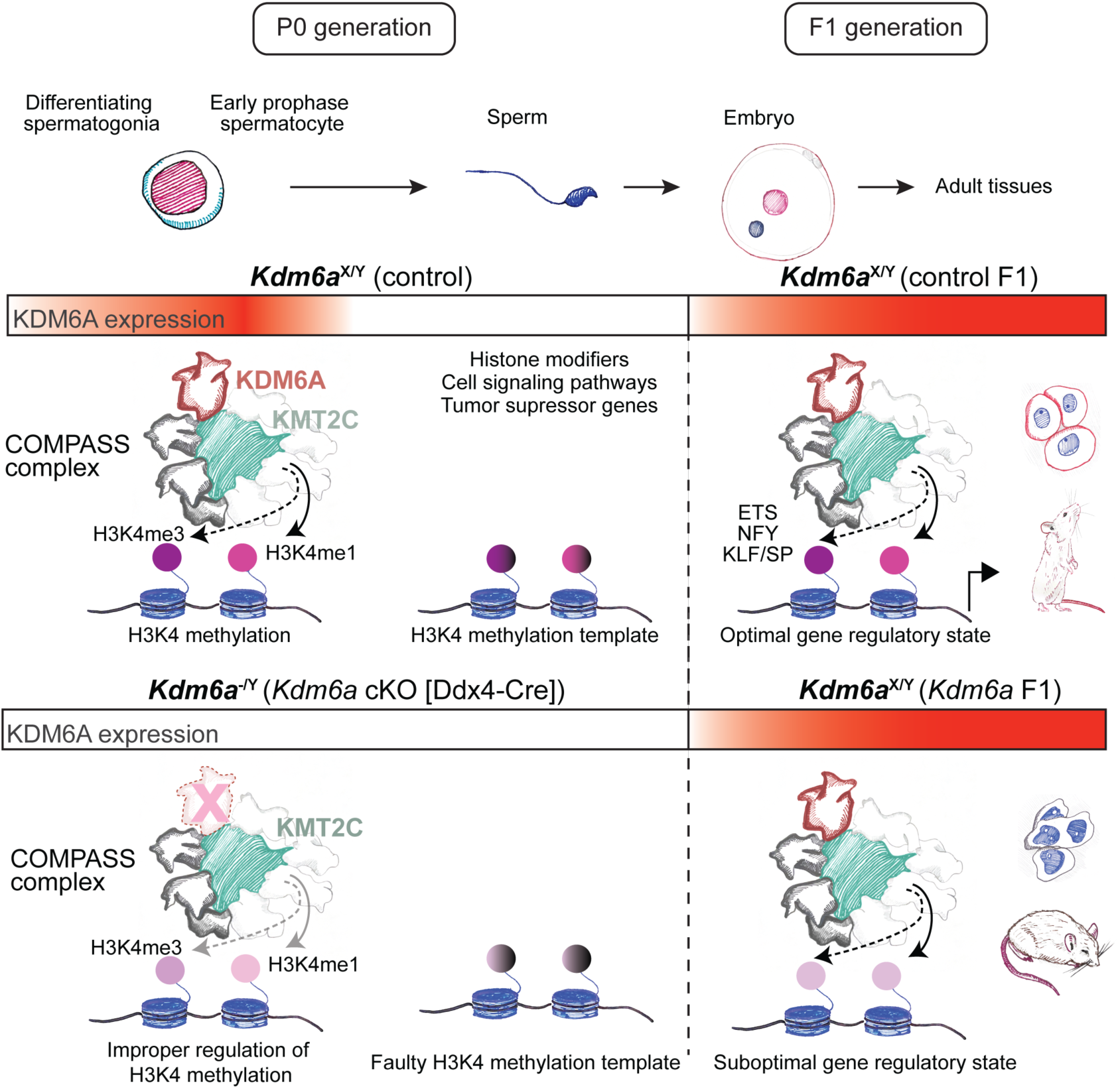
Model for epigenetic transmission of cancer susceptibility via dysregulation of KDM6A-COMPASS in the paternal germ line. **A,** KDM6A-COMPASS functions at the mitotic-meiotic transition in spermatogenesis to regulate the balance of H3K4 methylation at promoters, including tumor suppressors and histone modifiers. This activity influences H3K4 methylation state in sperm and may serve as a template for developing and mature tissues in the next generation. **B,** Loss of KDM6A disrupts COMPASS-mediated H3K4 methylation in differentiating spermatogonia leading to an abnormal histone methylation template in sperm and in the next generation. Suboptimum chromatin state at tumor suppressor genes or other cancer-relevant signaling pathways may predispose for malignant transformation.

A striking finding in our study is the strong enrichment of KDM6A at promoters in germ cells, compared to its preference for intergenic enhancers in other cell types including somatic cells of the testes. Most recent studies characterizing KMT2C and KDM6A have focused on their activity at enhancers (66). On the other hand, KMT2C has also been shown to regulate H3K4me1 and H3K4me3 at promoters in myoblasts, adult mouse brain, bladder carcinoma and leukemic cell lines (36, 67–69). In myoblasts, KMT2C-COMPASS deposits H3K4me1 at a subset of promoters to repress gene expression, and upregulation of these genes at differentiation correlates with loss of KMT2C, reduced H3K4me1/H3K4me3 ratio, and a switch to recruitment of SETD1A-COMPASS (69). Similarly, treatment of leukemic cells with a Menin-MLL1 inhibitor led to a switch at promoters from MLL1-Menin to KMT2C/D-KDM6A and a concomitant increase in H3K4me1 relative to H3K4me3. These findings indicate that multiple COMPASS family members can co-regulate a subset of promoters under different developmental contexts to balance H3K4me3 and H3K4me1 (70). Notably, while previous studies found at most one-third of KMT2C binding at promoters, we found that in germ cells promoters are the predominant site of KMT2C enrichment (69%). These results point to an unappreciated role for KMT2C at promoters that is especially prominent in male germ cells.

Another surprising finding was the absence of a verifiable interaction between KDM6A and the KMT2C paralog KMT2D, despite the similar biochemical activities of KMT2C and KMT2D and the well-established functional relationship between KDM6A and KMT2D in other tissues. This differential interaction is likely attributable to the coordinated expression between KDM6A and KMT2C during spermatogenesis. Interestingly, KMT2D is reported as a KDM6A partner in somatic tissues more frequently than KMT2C, and KDM6A and KMT2D mutations lead to similar clinical presentations in Kabuki syndrome while KMT2C mutations instead contribute to autism spectrum disorders. These findings suggest that KDM6A-KMT2D activity at enhancers is more widespread, while the preferential interaction of KDM6A with KMT2C may be a specialized germ cell function. Single and double conditional knockouts of *Kmt2c* and *Kmt2d* in the male germ line will be necessary to delineate their redundant and non-redundant roles in spermatogenesis, and to assess whether loss of KMT2C induces changes similar to those observed in the *Kdm6a* cKO germ line. Interestingly, our data also reveal partially nonredundant functions of KDM6A and its paralog KDM6B (71–73) in germ cells. While both *Kdm6a* and *Kdm6b* germline cKO mice are fertile and lack changes in H3K27me3, suggesting that they are redundant for fertility and H3K27me3 deposition (2, 31, 74), our results support a nonredundant function for KDM6A in H3K4 methylation during spermatogenesis.

While our data strongly support an impact of KDM6A on H3K4 methylation in sperm and on gene expression in the next generation, not all genes whose promoters had altered H3K4 methylation in *Kdm6a* cKO sperm had detectable gene expression changes in offspring. These results reinforce many previous observations of high variability in the correlation between epigenomic alterations in sperm and gene expression in offspring (75). Such variability is expected due to tissue-specific gene expression patterns, stochasticity in epigenetic transmission, and secondary alterations to the epigenome and transcriptome during development. Importantly however, altered expression of a single gene can be sufficient to elicit phenotypes in offspring (76). Since alterations to H3K4 methylation in *Kdm6a* cKO sperm preferentially occur at key tumor suppressor genes and cell signaling pathways, only a handful of misregulated genes may be sufficient to alter cancer risk. Furthermore, our observation of altered H3K4 methylation at *Kmt2c* and *Kmt2d* in the *Kdm6a* cKO paternal germ line, coupled with the downregulation of these genes in offspring, raises the possibility of a KDM6A–KMT2C positive feedback loop that crosses generations and may play a critical role in maintaining appropriate H3K4 methylation patterns. These findings amplify a growing literature implicating H3K4 methylation as a key element in intergenerational epigenetic transmission (14, 20, 21, 77–79).

In addition to COMPASS components, we identified several other KDM6A interacting proteins in male germ cells, including INO80C, SETD3, Mediator complex components, tRNA synthase complex components, and stress granule components. The INO80 complex is an essential meiotic factor in mouse spermatogenesis and was recently shown to interact with COMPASS and aid in regulating H3K4 methylation in *Arabidopsis* (43, 44). KDM6A may bridge the INO80 and COMPASS complexes, allowing cooperation between ATP-dependent remodeling and H3K4 methylation; a similar bridging role for KDM6A has been reported between the mitotic deacetylase complex MiDAC and KMT2C/D-COMPASS in mammalian cell lines (80). SETD3, another top interactor, has been primarily characterized as a methyltransferase of non-histone targets, although one study has described a role of SETD3 in H3K4/36 methylation in muscle differentiation, and SETD3 could also partner with KDM6A for H3K4 methylation during spermatogenesis. Meanwhile, alterations to sperm-borne tRNA fragments and mitochondrial tRNAs have proven to be key mediators of epigenetic transmission in sperm (16, 17, 81); interactions with tRNA synthase components suggest the possibility that KDM6A may contribute to regulating the composition of this tRNA payload. Finally, KDM6A was recently found to re-localize to the cytoplasm under stress conditions in vitro and associates with stress granules via interactions with G3BP1, also identified as a KDM6A interactor in our data. Regulation of stress granule disassembly by KDM6A was found to be important for suppressing cancer. It is possible that the *Kdm6a* cKO male germ line are less resilient to stress due to a loss of this activity, amplifying accumulation of more genetic and epigenetic changes.

Mutations in COMPASS histone methyltransferases are among the most frequent alterations in human cancers. Double deletion of *Kmt2c* and *Kmt2d* in mouse urothelium or loss of *Kdm6a* in mouse hematopoietic stem cells appears to prime cells for malignant transformation in these tissues via alterations to H3K4 methylation, reminiscent of the intergenerational changes we observed (3, 82). Thus, the anti-oncogenic epigenetic functions of KDM6A, KMT2C, and the COMPASS complex that are important in somatic tissues may also operate transgenerationally. Our data indicate that loss of KDM6A-KMT2C COMPASS activity in germ cells establishes tumor-promoting epigenetic alterations that are transmitted to the next generation, lowering the barrier to cancer. Indeed, we observed that altered H3K4me3 in *Kdm6a* cKO sperm occurs preferentially at promoters of genes encoding tumor suppressors and components of key cell signaling pathways implicated in tumorigenesis. However, whether changes in H3K4 methylation are directly transmitted to the next generation or instead undergo epigenetic reprogramming in the embryo and are re-established at the same loci during development remains an open question. Altogether, our data support H3K4 methylation in the male germ line as a key mediator of epigenetic inheritance and highlight altered regulation of this histone modification as a contributing mechanism for intergenerational cancer susceptibility.

## Materials & Methods

### Data availability

All sequencing data generated for this study is available from the NCBI Gene Expression Omnibus under accession number GSE334525. Proteomics data is available at the Proteome Xchange PRIDE database under accession number. Public datasets used are listed in **Supplementary Table S5.**

### Animals

All mice were maintained on a C57BL/6J genetic background. To obtain *Kdm6a* cKO males, *Kdm6a*^flox/flox^ (*Kdm6a^tmc1(EUCOMM)Jae^*) females were mated with mice carrying one copy of the *Ddx4*-Cre transgene (B6-*Ddx4^tm1.1(cre/mOrange)Dcp^*) (83, 84) in which Cre is expressed specifically in germ cells beginning at embryonic day 15.5. *Kdm6a* F1 embryos were generated by crossing male *Kdm6a* cKO mice with wild type females. These studies were approved by the Yale University Institutional Animal Care and Use Committee under protocol 2023-20169. All mice used in these studies were maintained and euthanized according to the principles and procedures described in the National Institutes of Health guide for the care and use of laboratory animals.

### Antibodies

Antibodies used in this study are listed in **Supplementary Table S6.**

### ChIP-qPCR primers

ChIP-qPCR primers used in this study are listed in the **Supplementary Table S7.**

### Immunostaining

Adult mouse testes were punctured with an insulin needle and immersed in 10 volumes of Hartman’s fixative (or 4 % paraformaldehyde for vimentin staining) for 1 h at room temperature. Samples were then bisected and fixed overnight at 4 °C with agitation. Fixed testes were dehydrated, cleared, and embedded in paraffin. Sections were dewaxed in three changes of xylene and rehydrated through two changes each of 100%, 95%, and 70% ethanol, followed by distilled water.

Antigen retrieval was performed by boiling sections in citrate buffer (Abcam, AB93678) for 11 min, cooling to 50 °C on ice, and then reheating for an additional 3 min. After cooling to room temperature, slides were rinsed in PBS and incubated in 0.3% Sudan Black B for 1 h at room temperature. Sections were then washed and blocked in 5% bovine serum albumin (BSA)/PBS for 1 h at room temperature.

Primary antibodies diluted in 1% BSA/PBS with applied overnight at 4 °C. After PBS washes, Alexa Fluor–conjugated secondary antibodies diluted in 1% BSA/PBS were applied for 1 h at room temperature. Sections were washed, counterstained with Hoechst for 5 min, rinsed in PBS, and coverslipped with 90% glycerol/PBS. Images were acquired using a Zeiss epifluorescence microscope.

### Whole cell protein extraction

Cultured cells (1 × 10⁶), KIT⁺-sorted cells (1 × 10⁵), or seminiferous tubules (10 mg) were homogenized in RIPA buffer supplemented with 0.9% SDS, protease inhibitor cocktail (Roche, 11836170001), and 5 mM sodium butyrate (Sigma-Aldrich, 303410). Extraction of proteins from sperm was performed by resuspending the sperm cell pellet obtained from two cauda epididymides as described above in 100 uL pre-boiled SDS lysis buffer containing 10 mM Tris-HCl (pH 8.0), 1 mM sodium orthovanadate, and 1% SDS. These samples were then heated at 100 °C for 10 min and cooled to room temperature. Both SDS and RIPA lysates were sheared by passage through an insulin syringe needle to reduce viscosity, followed by centrifugation at 13,600 × g for 30 min. Supernatants were transferred to fresh tubes and total protein concentration was determined using a BCA assay (Thermo, 23225). Protein concentrations were equalized across samples before denaturing with 4x Laemmli buffer containing β-mercaptoethanol at 95 °C for 5 min.

### Western blotting

IP protein samples were resolved on Tris–acetate gels (Invitrogen, EA0375BOX), while whole-cell lysates were separated on 4–20% Tris–glycine gels (Bio-Rad, #4568093) at 180 V for 50 min. Proteins were then wet transferred onto 0.45 μm nitrocellulose membranes (BioRad) in NuPAGE buffer (for Tris–acetate gels, Invitrogen, #NP00061) or Towbin buffer (for Tris–glycine gels) at 350 mA for 70 min. Membranes were blocked for 1 h in 5% nonfat dry milk (Research Products International, M17200) in TBST, followed by overnight incubation at 4 °C with primary antibodies diluted in 5% BSA/TBST. After washing with TBST, membranes were incubated for 1 h at room temperature with Horseradish peroxidase-conjugated secondary antibodies diluted in blocking solution. Following additional TBST washes, membranes were incubated with chemiluminescent substrate (Thermo, #34580) for 5 min and exposed to X-ray film.

### GSK-J4 treatment of immortalized testicular cell lines

GC1-spg and GC2-spd cells were cultured to 80% confluency in a 6 well dish and treated with GSK-J4 (MedChemExpress, HY-15648B) or DMSO for 24 h at 37 °C with 5 % CO_2_. Cells were trypsinized, washed in PBS, then pelleted before extracting proteins with RIPA as described above.

Testes were dissociated as described above and plated in a 6 well dish containing DMEM/F12 (1:1) (Gibco, 1130-032) supplemented with 10% FBS. Cells were then immediately treated with DMSO or GSK-J4 and incubated for 24 h at 37 °C with 5 % CO_2_. Adherent cells were trypsinized and pelleted along with the non-adherent cells before extracting proteins with RIPA described above.

### Generation of transgenic GC1-spgs

Lentiviral particles were generated from HEK 293T cells cultured in a 6 cm dish using Lipofectamine 3000 (Thermo, L3000001), 1ug PAX2, 0.5ug VSV-G and 1.5 ug of KDM6A DNA expression constructs (Y. Soto-Feliciano). Viral supernatants were harvested at 24 h and 72 h post-transfection and centrifuged at 2,000 rpm for 10 mins. The clarified supernatant (8 ml) was syringe filtered through a 0.45 μm filter and then mixed with 2 ml PEG-it virus precipitation solution (System Biosciences, LV810A-1) followed by incubation overnight at 4 °C. The precipitated viral supernatant was centrifuge at 1,500 x g for 30 mins at 4 °C. The resulting viral pellet was resuspended in 100 μl of cold DPBS and used immediately to reverse transduce 100,000 GC1-spgs in a 6 well format.

Immortalized testis cultures expressing shScramble or sh*Kdm6a* were generated in a previous study (31).

All cell lines were free of mycoplasma contamination.

### Co-immunoprecipitation of nuclear proteins

Seminiferous tubules (30 mg) isolated from an adult mouse testis or a 150 mm dish of confluent GC1-spgs were used per immunoprecipitation (IP) experiment. Samples were processed using a Nuclear complex Co-IP kit (Active Motif, #54001) according to the manufacturer’s instructions, with low-stringency IP buffer used throughout.

For mass spectrometry, samples were incubated with 50 ul of washed Dynabeads Protein G (Thermo Fisher Scientific, #10004D) pre-crosslinked to primary antibody using 5 mM bis(sulfosuccinimidyl) suberate (Thermo, #21580). For Western blot validation, samples were first incubated with primary antibody overnight at 4 °C, followed by incubation with Dynabeads Protein G for 1h at 4 °C.

Beads were washed three times in 1 ml of IP wash buffer containing 1 mg/ml BSA, followed an additional three washes in IP wash buffer without BSA. The washed beads were resuspended in 20 ul of LDS sample buffer for Western blotting, or Laemmli buffer for mass spectrometry analysis, and heated at 70 °C in the presence of reducing agent for 10 mins.

### Mass spectrometry and analysis

For analysis by mass spectrometry, proteins were cleaned up/digested using S-Trap micro columns (ProtiFi, #C02-micro-40) following the manufacturer’s protocol with minor adjustments. Briefly, samples were diluted with 50 uL of 10% SDS and acidified with trifluoracetic acid (TFA) to a final concentration of 1%. Six volumes of S-Trap binding buffer (90% methanol/100 mM TEAB, pH = 8.5) were added to each sample and samples were loaded onto S-Trap columns via centrifugation. Proteins were alkylated on-column by incubating with 200 uL of 10 mM iodoacetamide (Sigma-Aldrich, #I1149) in binding buffer for 10 min at room temperature in the dark. The iodoacetamide solution was spun through the column and the alkylation step was repeated once. Columns were washed three times with 50/50 methanol/chloroform followed by four times with binding buffer, then samples were digested at 37 °C via overnight incubation with 0.5 ug trypsin (Promega, cat. #V5113) in 50 mM ammonium bicarbonate buffer. Peptides were eluted via centrifugation and columns were washed successively with 50 mM ammonium bicarbonate, 0.2% TFA, and 50/50 water/acetonitrile (ACN). Peptides were dried via vacuum centrifugation and resuspended in 25 ul of 0.1% formic acid (FA) in water.

LC-MS/MS data were acquired on a Thermo Scientific Exploris 480 mass spectrometer coupled to a Thermo Scientific Vanquish Neo UHPLC system. Peptides (5 uL) were loaded onto a PepMap Neo C18 trap cartridge (100 Å, 5 µm, 300 μm × 5 mm) (Thermo Scientific, cat. #174500) at a maximum pressure of 800 bar and separated using an Aurora Ultimate C18 analytical column (120 Å, 1.7 μm, 75 μm × 250 mm) (IonOpticks, cat. #AUR3-25075C18-TS) (50 °C). The compositions of mobile phases A and B were 0.1% FA in water and 0.1% FA in 80% ACN, respectively. The peptides were eluted at a flow rate of 300 nL/min with a gradient starting at 1% B, increasing to 4% B over 4 min, then to 25% B over 84 min and 40% B over 32 min. The gradient was then ramped to 450 nL/min and 99% B over 1 min and held for 10 min before equilibrating to starting conditions. Precursor MS1 scans (profile) were collected from 350-1500 m/z at a resolution of 120,000. The AGC target was set to 3 × 106 and the maximum injection time was 50 ms. Data-dependent MS2 scans (centroid) were collected in top 20 fashion for precursors with charge states 2-6. The isolation window was set to 1.4 m/z and HCD fragmentation was employed with a collision energy setting of 28%. MS2 spectra were collected starting at 120 m/z with a resolution of 22,500. The AGC target was set to 1 × 105 and the maximum injection time was 50 ms. Dynamic exclusion was enabled using automatic settings and at least four blank injections were carried out in between sample injections to minimize carryover.

Protein identification was performed in Proteome Discoverer (Thermo, version 3.0.1.27) using standard processing and consensus workflows. Briefly, precursor masses were recalibrated before data were searched against the SwissProt Mus musculus database (downloaded October 2024) and a database of common contaminant proteins. The CHIMERYS 1.0 search engine (85) was employed using the INFERYS 2.1 prediction model (86). Trypsin was selected as the enzyme, up to two missed cleavages were allowed, and the fragment ion mass tolerance was set to 0.02 Da. Peptides with lengths between 7-30 amino acids and charge states +1-6 were allowed. Oxidation of methionine was set as a variable modification and carbamidomethylation of cysteine was set as a fixed modification. False discovery rates (FDRs) were estimated using Percolator (87). Peptide- and protein-level FDR thresholds were set to 1% and only master proteins with at least two protein unique peptides were considered for downstream analysis. KDM6A protein interactors were qualitatively identified by those consistently detected across all KDM6A IP replicates but not in IgG control samples.

### MNase ChIP in sperm

Cauda epididymides from adult mice were cut into Donner’s medium (135 mM NaCl, 5 mM KCl, 1 mM MgSO4, 2 mM CaCl2, 30 mM HEPES, 25mM NaHCO3, 2% BSA, 1mM sodium pyruvate, 0.32% sodium DL-lactate) and incubated at 37 °C for 1 h. The resulting sperm suspension was filtered through a 40 um nylon mesh and centrifuged at 2,500 x g for 8 min at 4 °C.

The pellet was washed once with PBS, then with 0.45% NaCl, and once again with PBS with centrifugation between each step as described above. Cells were then resuspended in 1 mL PBS and treated with 100 uL RDD buffer and 25 uL DNase (Qiagen #79254) to remove extracellular DNA. Samples were centrifuged at 2,000 x g for 5 min at room temperature, resuspended in 1 mL PBS, flash frozen in liquid nitrogen, and stored at -80 °C. Each ChIP experiment used sperm pooled from three mice.

Buffers were prepared as follows. Buffer 1 contained 15 mM Tris-HCL, 60 mM KCl, 5 mM MgCl2, and 0.1 mM EGTA in water. MNase buffer contained 85 mM Tris-HCl, 3 mM MgCl2, and 2 mM CaCl2 in water. Combined Buffer contained equal parts Buffer 1 and MNase buffer with 0.3M sucrose. Wash buffer A contained 50 mM Tris-HCl, 10 mM EDTA, and 75 mM NaCl in water. Wash Buffer B contained 50 mM Tris-HCl, 10 mM EDTA, and 125 mM NaCl in water. Elution buffer contained of 0.1M NaHCO3, 0.2% SDS, 5 mM DTT, 10mM Tris-HCl, and 1mM EDTA in water.

Dynabeads Protein G were washed twice in 10 mM Tris-HCl with 1 mM EDTA, followed by two additional washes in combined buffer. Beads were then incubated for 3 h at 4 °C in combined buffer supplemented with 1 mg/mL BSA and 100 μL of tRNA solution (Sigma Aldrich #R8508). After incubation, beads were washed once with combined buffer, resuspended in combined buffer, and kept at 4 °C until use.

Frozen sperm samples were thawed at room temperature, pelleted by centrifugation, and resuspended in 1 mL PBS. After addition of 50 µL of 1M DTT, samples were incubated at room temperature for 2 h. Subsequently, 120 µL of 1M N-Ethylmaleimide (Millipore #128-53-0) was added, and samples were incubated for an additional 30 min at room temperature. Samples were washed once with PBS and resuspended in 300 µL of Buffer 1 supplemented with 0.3M sucrose and 1mM DTT, followed by the addition of 300 µL Buffer 1 with 0.3 M sucrose, 1 mM DTT, 0.5% NP-40 (Sigma-Aldrich #NP40S), and 1 % sodium deoxycholate (Sigma-Aldrich, #D6750).

Samples were incubated on ice for 10 min and then divided into 100 uL aliquots. Each aliquot was mixed with 100 uL MNase buffer containing 0.3 M sucrose and 400 U/mL micrococcal nuclease S7 (Roche, #10107921001) and incubated for 5 min at 37 °C. The reaction was stopped by adding 2 uL of 0.5 M EDTA. Aliquots were placed on ice for 5 min and centrifuged at 16,300 x g for 10 min. Supernatants of each individual sample were recombined, treated with 65 uL of 20x protease inhibitor (Roche #11836170001), and precleared by incubation with 50 uL of washed beads 1 h at 4 °C with rotation.

Following centrifugation, supernatants were transferred to fresh microcentrifuge tubes, and 100 uL was reserved as input. Primary antibody was added to the remaining samples and incubated overnight at 4 °C. The following morning, samples were incubated with 50 μL of washed beads for 6 h at 4 °C with rotation. Bead-bound complexes were washed once with Wash Buffer A and twice with Wash Buffer B

Chromatin was eluted by incubating beads with elution buffer for 15 min at 65 °C, and eluates were combined. Both immunoprecipitated and input chromatin samples were treated with 6 μL RNase A (Sigma-Aldrich, #70856) for 30 min at 37 °C, followed by incubation with 6 uL proteinase K (Thermo, #AM2548) overnight at 55 °C. DNA was purified the next morning using the ChIP DNA Clean & Concentrator kit (Zymo, #D5201).

#### Crosslinked ChIP in testis

Testes from adult mice were decapsulated, and 30 mg of tubules were processed for ChIP-seq following the protocol and buffer formulations described by Sullivan and Santos (88), with minor modifications. Tubules were minced in cold DPBS and titurated by repeated pipetting until no tissue fragments larger than 0.5 mm remained. The resulting cell suspension was transferred to a 15 ml tube containing 5 ml of DPBS at room temperature, and 40 μl of 300 mM DSG (Thermo, #20593) was added. Samples were incubated for 30 min at room temperature with rotation.

Formaldehyde (Thermo, #28906) was then added to a final concentration of 1% and incubated for 10 min at room temperature with rotation. Crosslinking was quenched by the adding glycine to a final concentration of 125 mM, followed by incubation for 5 min at room temperature.

Fixed cells were washed twice in 6 ml DPBS containing 0.5% BSA by centrifugation at 760 x g for 5 min at 4 °C, then resuspended in 1 ml DPBS and transferred to a 1.5 ml tube.

Cells were pelleted at 380 x g for 5 min at 4 °C and resuspended in 300 ul lysis buffer supplemented with protease inhibitors (Roche). Samples were transferred to 1.5 ml sonication tubes, and chromatin was sheared using a Bioruptor Pico device (Diagenode) for 10 cycles of 30 secs on/off at 4 °C.

Sheared chromatin was diluted with 1 ml dilution buffer supplemented with protease inhibitors (Roche) and clarified by centrifugation for 13,600 x g for 30 min at 4 °C. The supernatant was transferred to fresh 1.5 ml tubes, and 1% of the sample was reserved as input. Samples were incubated overnight with primary antibody at 4 °C with rotation. The following morning, 20 ul of pre-washed Dynabeads Protein G (Thermo) were added and incubated for 3 h at 4 °C with rotation. Beads were then washed, and immunocomplexes were eluted, followed by reverse crosslinking as previously described. DNA was purified using the ChIP DNA Clean & Concentrator kit (Zymo).

#### ChIP-qPCR

For ChIP-qPCR, each reaction contained 10 ul PowerUp SYBR Green Master Mix (Applied Biosystems, #A25742), 1 ul of 10 μM primer mix, 1 ul DNA and 8 ul nuclease-free water. qPCR was performed in 96-well plates using a QuantStudio 3 system (Applied Biosystem).

### ChIP-seq library preparation

ChIP-seq libraries were prepared using the Watchmaker DNA Library Prep Kit (Watchmaker Genomics, Part# 7K0101-096) following the manufacturer protocol. Adapter-ligated DNA fragments were PCR amplified (8 -10 cycles) using custom-made primers. During PCR, a unique 10 base index was inserted at both ends of each DNA fragment. Size of the final library construct was determined on Agilent Tape Station system and quantification is performed by qPCR using the Kapa Library Quantification Kit. Samples with a yield of ≥0.5 ng/ul were used for sequencing.

### CUT&Tag in differentiating spermatogonia

Differentiating spermatogonia were purified and CUT&Tag libraries were prepared as previously described (89), except that 0.05% trypsin-EDTA was used for cellular dissociation instead of Accutase. Approximately one-hundred thousand KIT+ cells were used for each CUT&Tag reaction using the CUT&Tag-IT assay kit (Active Motif, 53160).

### Pre-implantation embryo RNA-seq

Female mice (B6D2F1) were superovulated via intraperitoneal injection of 7 IU pregnant mare serum gonadotropin (PMSG), followed 48 hours later by an injection of 5 IU human chorionic gonadotropin (hCG) and then mated with wildtype or *Kdm6a* cKO male mice. Zygotes were harvested from the oviducts 16–20 hours after hCG injection and cultured in KSOM medium under mineral oil at 37°C in atmosphere of 5% CO_2_ until the desired stage. Thirty embryos for each sample were collected in 100 ul of lysis buffer. Two replicates of each embryonic stage were collected. RNA was extracted according to the manufacturer’s protocol using the RNeasy Micro kit (Qiagen) and RNA-seq libraries were prepared using SMARTer Stranded Total RNA-Seq kit v3 – Pico Input Mammalian (Takara, #634486) according to manufacturer’s instructions.

### Processing and analysis of sequencing datasets

ChIP-seq and CUT&Tag libraries were sequenced on an Illumina NovaSeq platform using 150 bp paired-end reads, at a depth of 30 million reads per sample. Filtering and adapter trimming for ChIP-seq and CUT&Tag reads was performed using the default settings of TrimGalore (90). Filtered ChIP-seq reads were aligned to the mouse (mm10) reference genome with BowTie2 using the ‘end-to-end’ mode and ‘fast’ option and filtered CUT&Tag reads were aligned using the ‘local’ mode and ‘very-sensitive’ option (91). The resulting SAM files were converted to BAM files and coordinate sorted with SAMtools (92). BigWig files were generated using the bamCompare (ChIP-seq datasets) or bamCoverage (CUT&Tag datasets) modules in deepTools (93) and visualized on the Integrative Genomics Viewer (94). Narrow peaks (H3K4me3, KDM6A and KMT2C) and broad peaks (H3K4me1) were called using MACS2 with the appropriate input as background (95). Heatmaps of average genomic signal intensity were generated using the computeMatrix and plotHeatmap modules in deepTools. Differential analysis (edgeR) and quality control analysis of ChIP-seq datasets were performed using DiffBind version 3.14.0 (96). Descriptive stats of ChIP-seq and CUT&Tag datasets was performed using ChIPseeker (97). Gene ontology analysis was performed using EnrichGO (39).

RNA-seq libraries were sequenced on an Illumina NovaSeq to generate paired-end, 150 base pair read libraries at an average depth of 35 million read pairs/library. RNA-seq reads were pseudoaligned to the mouse (mm39) reference transcriptome using Kallisto and differentially expressed genes between groups were identified using DESeq2 with adjusted p-value 0.10 as a significance threshold.

## Acknowledgements

We helpful discussions with members of the Lesch lab. We thank the Yale Center for Genome Analysis for library preparation and high-throughput sequencing. We thank the Keck Mass Spectrometry & Proteomics Resource at the Yale School of Medicine for providing the necessary LC-MS/MS instruments and accompanying software funded in part by the Yale School of Medicine and by the Office of the Director, National Institutes of Health (S10OD02365101A1, S10OD019967, and S10OD018034). The funders had no role in study design, data collection and analysis, decision to publish, or preparation of this manuscript.

B.W.W. was supported by a Hope Funds for Cancer Research Postdoctoral Fellowship. This work was supported by the National Institute of Child Health and Human Development (R01HD098128 and R21HD110843 to B.J.L.), the National Cancer Institute (R21CA288677), an ACS-IRG pilot award from the Yale Cancer Center, and the Searle Scholars Program. Bluma Lesch, M.D., Ph.D. was supported by a Discovery Boost Grant, DBG-23-1150177-01-DMC, Grant DOI #10.53354/ACS.DBG-23-1150177-01-DMC.pc.gr.175431, from the American Cancer Society. B.J.L. is a Pew Scholar, supported by the Pew Charitable Trust.

## Author contributions

Conceptualization: BWW, BJL

Investigation: BWW, RAH, HY, SK, JT, KH

Formal analysis: BWW, KH

Resources: ZL, YS-F, BJL

Validation: BWW

Visualization: BWW

Writing – original draft: BWW

Writing – review & editing: BJL

Funding acquisition: BJL

