## Supplemental Figures for "Paternal regulation of H3K4 methylation supports tumor suppressor networks in mammals intergenerationally"

### Figure S1

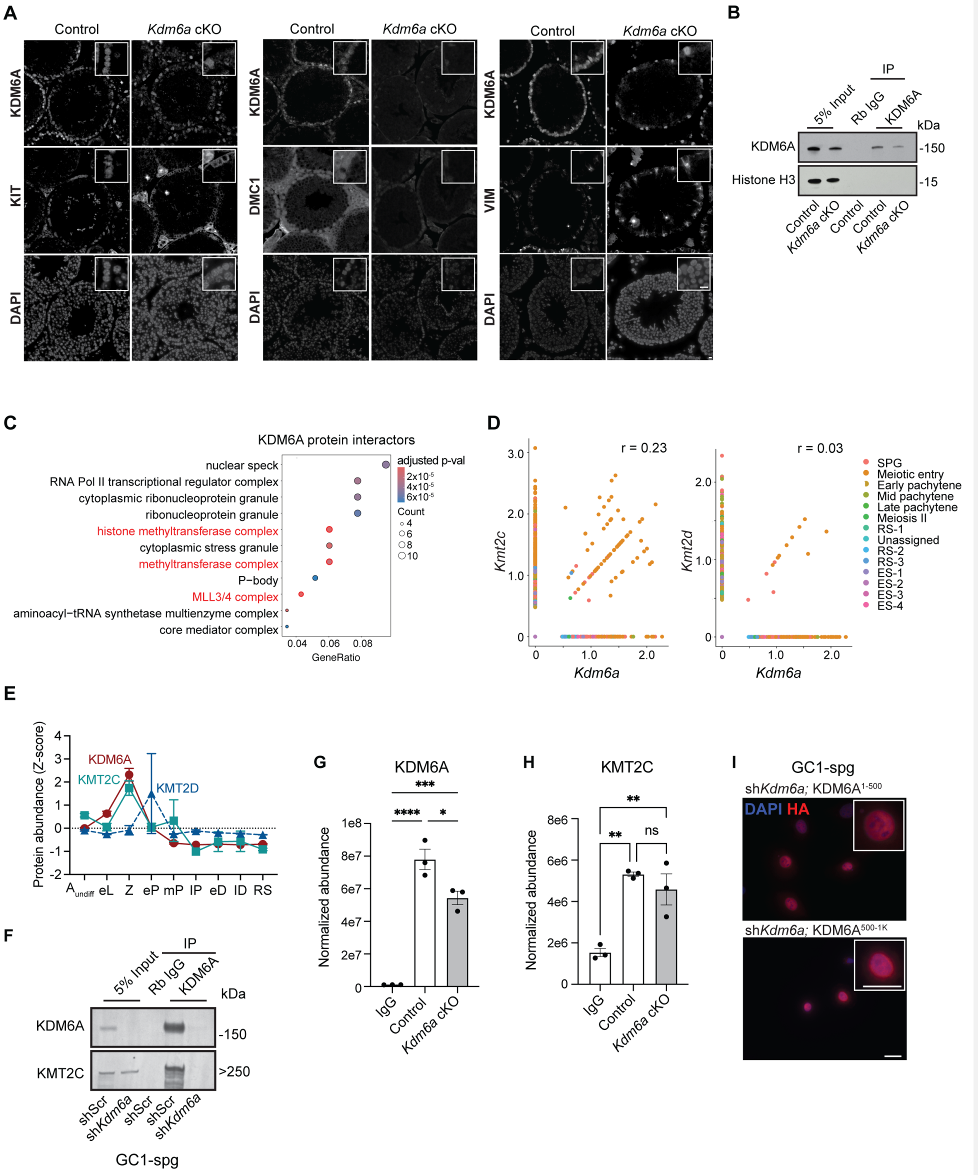
A
Control
Kdm6a cKO
KDM6A
KIT
DAPI
DMC1
VIM
B
5% Input
IP
Rb IgG
KDM6A
kDa
KDM6A
-150
Histone H3
-15
Control
Kdm6a cKO
C
KDM6A protein interactors
nuclear speck
RNA Pol II transcriptional regulator complex
cytoplasmic ribonucleoprotein granule
ribonucleoprotein granule
histone methyltransferase complex
cytoplasmic stress granule
methyltransferase complex
P−body
MLL3/4 complex
aminoacyl−tRNA synthetase multienzyme complex
core mediator complex
adjusted p-val
Count
GeneRatio
D
r = 0.23
r = 0.03
Kmt2c
Kmt2d
Kdm6a
SPG
Meiotic entry
Early pachytene
Mid pachytene
Late pachytene
Meiosis II
RS-1
Unassigned
RS-2
RS-3
ES-1
ES-2
ES-3
ES-4
E
Protein abundance (Z-score)
KDM6A
KMT2C
KMT2D
eL
Z
eP
mP
IP
eD
ID
RS
F
5% Input
IP
Rb IgG
KDM6A
kDa
KDM6A
-150
KMT2C
>250
shScr
shKdm6a
GC1-spg
G
KDM6A
Normalized abundance
IgG
Control
Kdm6a cKO
H
KMT2C
Normalized abundance
ns
I
GC1-spg
shKdm6a; KDM6A1-500
DAPI HA
shKdm6a; KDM6A500-1K

**Fig. S1. Analysis of KDM6A protein interactions in mouse testis. A,** Individual epifluorescence channels (DAPI, 488 nm, and 594 nm) from the immunofluorescence images depicted in **Fig. 1A**. **B,** Representative Western blot for KDM6A in whole testis samples from control and *Kdm6a* cKO mice following immunopreciptation of endogenous KDM6A using anti-KDM6A crosslinked to Dynabeads Protein G. Rb IgG, Rabbit IgG. **C,** Top ranked Cellular Component gene ontology terms for KDM6A protein interactors identified in adult mouse testis. Terms related to COMPASS are highlighted in red. **D,** Correlation between expression levels of *Kdm6a* and *Kmt2c* (left) or *Kmt2d* (right) transcripts within the same cell from testis single cell RNA-seq (*26*). SPG, spermatogonia; SPC, spermatocytes; RS, round spermatids; ES, elongating spermatids. r = Pearson correlation coefficient. **E,** Protein abundances for KDM6A, KMT2C, and KMT2D estimated from mass spectrometry of sorted spermatogenic cells at different stages of development: undifferentiated spermatogonia type A ($A_{undiff}$), early leptotene (eL), zygotene (Z), early pachytene (eP), mid pachytene (mP), late pachytene (lP), early diplotene (eD), late diplotene (lD), and round spermatids (RS). Error bars, s.e.m. Data from Fang et al. 2021 (*95*). **F,** Western blot for KDM6A and KMT2C in samples from GC1-spg cells expressing shScramble (shScr) or sh*Kdm6a* after immunoprecipitating endogenous KDM6A. Rb IgG, Rabbit IgG. **G,** Normalized abundance of KDM6A protein measured by mass spectrometry in testis samples from control and *Kdm6a* cKO mice after immunoprecipitation with IgG or anti-KDM6A. Points represent individual replicates and bars indicate mean ± SEM. Statistical significance was determined using a one-way ANOVA followed by Tukey's multiple comparisons test. *p < 0.05, **p < 0.01, ***p < 0.001, ****p < 0.0001. **H,** Normalized abundance of KMT2C protein measured by mass spectrometry in testis samples from control and *Kdm6a* cKO mice after immunoprecipitation with IgG or anti-KDM6A. Points represent individual replicates and bars indicate mean ± SEM. Statistical significance was determined using a one-way ANOVA followed by Tukey's multiple comparisons test. ns = not significant, *p < 0.05, **p < 0.01, ***p < 0.001, ****p < 0.0001. **I,** Immunofluorescence detection of HA-tag (red) in GC1-spg cells expressing sh*Kdm6a* and the indicated HA-tagged, truncated KDM6A protein. Scale bar = 10μm.

Commented [MOU1]: Add Fang et al. reference

**Figure S2**

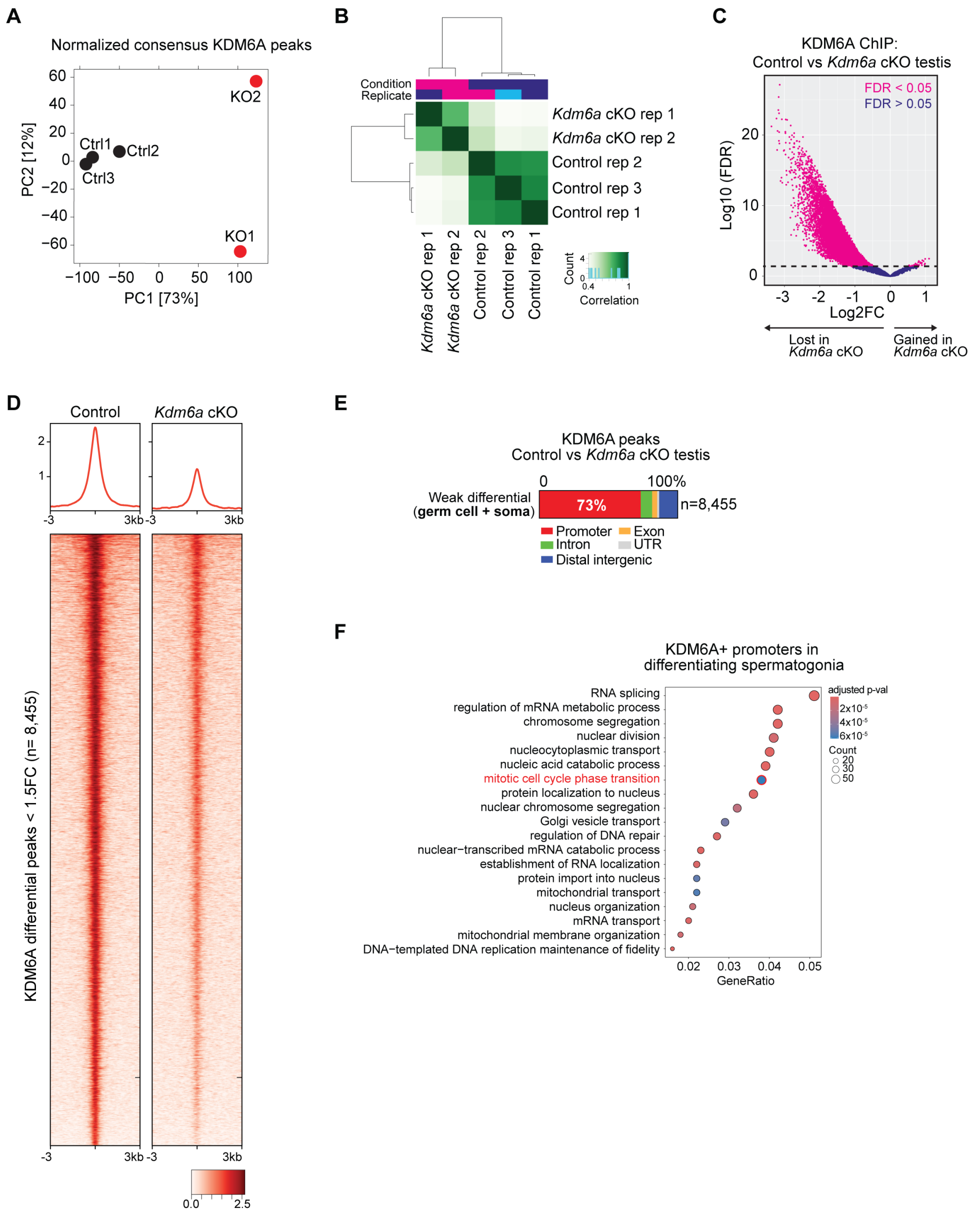
A
Normalized consensus KDM6A peaks
KO2
Ctrl1
Ctrl2
Ctrl3
KO1
PC2 [12%]
PC1 [73%]
B
Condition
Replicate
Kdm6a cKO rep 1
Kdm6a cKO rep 2
Control rep 2
Control rep 3
Control rep 1
Count
Correlation
C
KDM6A ChIP:
Control vs Kdm6a cKO testis
FDR < 0.05
FDR > 0.05
Log10 (FDR)
Log2FC
Lost in Kdm6a cKO
Gained in Kdm6a cKO
D
Control
Kdm6a cKO
KDM6A differential peaks < 1.5FC (n= 8,455)
E
KDM6A peaks
Control vs Kdm6a cKO testis
Weak differential (germ cell + soma)
73%
n=8,455
Promoter
Exon
Intron
UTR
Distal intergenic
F
KDM6A+ promoters in differentiating spermatogonia
RNA splicing
regulation of mRNA metabolic process
chromosome segregation
nuclear division
nucleocytoplasmic transport
nucleic acid catabolic process
mitotic cell cycle phase transition
protein localization to nucleus
nuclear chromosome segregation
Golgi vesicle transport
regulation of DNA repair
nuclear−transcribed mRNA catabolic process
establishment of RNA localization
protein import into nucleus
mitochondrial transport
nucleus organization
mRNA transport
mitochondrial membrane organization
DNA−templated DNA replication maintenance of fidelity
GeneRatio
adjusted p-val
Count

**Fig. S2. KDM6A ChIP-seq in control and *Kdm6a* cKO mouse testis.**

**A,** Principal component analysis of KDM6A ChIP-seq datasets using consensus KDM6A peaks from samples of control (ctrl) and *Kdm6a* cKO (KO) mouse testis. **B,** Correlation across replicates of KDM6A ChIP-seq datasets from control and *Kdm6a* cKO mouse testis samples. **C,** Differential binding analysis of KDM6A (DiffBind) showing differential (pink) and non-differential (purple) peaks between control and *Kdm6a* cKO ChIP-seq datasets. FDR = False discovery rate. **D,** RPKM-normalized signal for KDM6A ChIP-seq centered on the summits of KDM6A peaks exhibiting intermediate reductions (< 1.5 FC) in *Kdm6a* cKO testes relative to control as determined by DiffBind, inferred as binding events common between the testicular soma and germline. **E,** Distribution of genomic features for intermediate KDM6A peaks shown in (**D**). **F,** Top-ranked Biological Process gene ontology terms enriched among KDM6A peaks detected in differentiating spermatogonia. The 'mitotic cell cycle phase transition' term is highlighted in red.

**Figure S3**

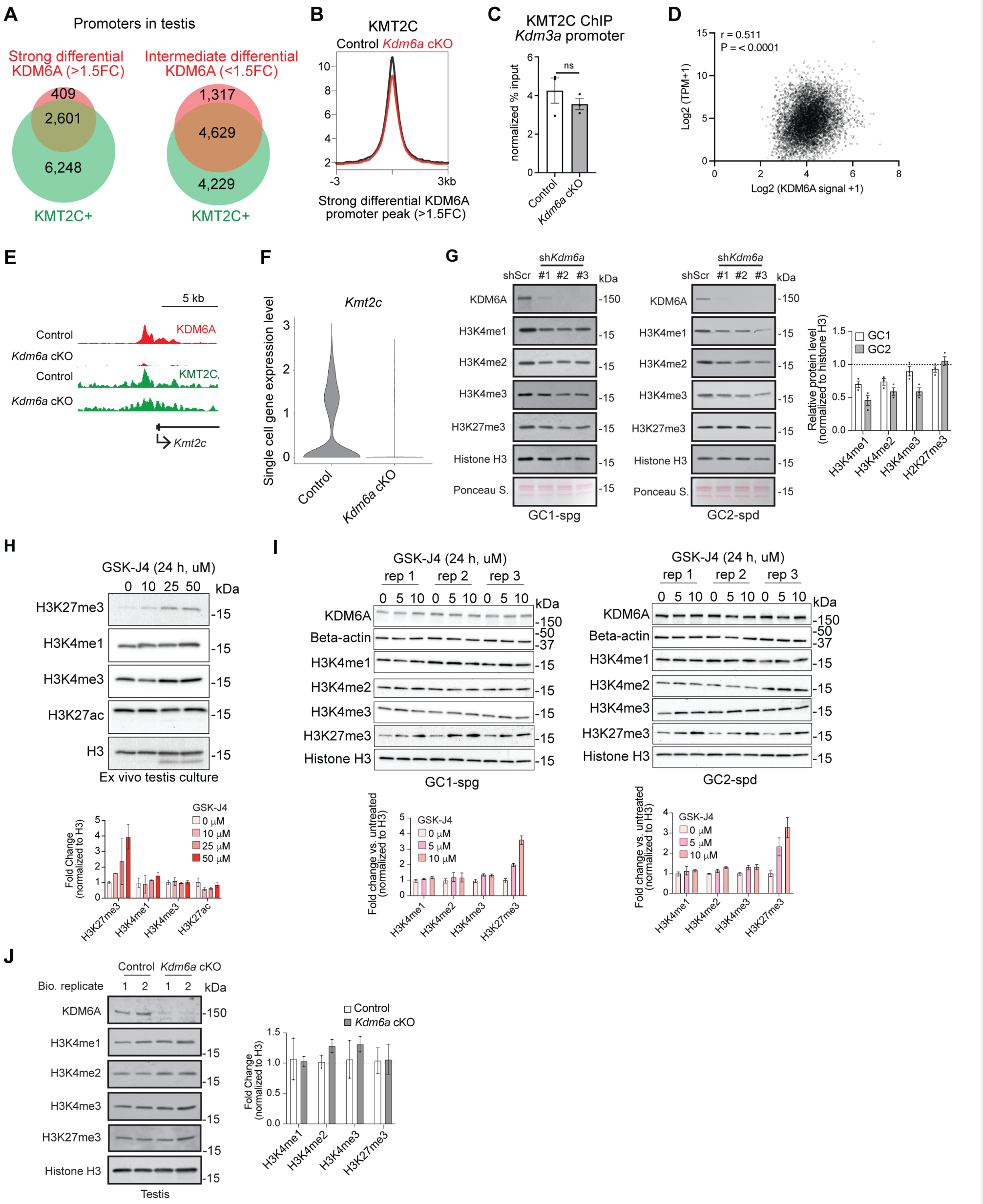
A
Promoters in testis
Strong differential KDM6A (>1.5FC)
409
2,601
6,248
KMT2C+
Intermediate differential KDM6A (<1.5FC)
1,317
4,629
4,229
KMT2C+
B
KMT2C
Control Kdm6a cKO
Strong differential KDM6A promoter peak (>1.5FC)
C
KMT2C ChIP
Kdm3a promoter
normalized % input
ns
Control
Kdm6a cKO
D
r = 0.511
P = < 0.0001
Log2 (TPM+1)
Log2 (KDM6A signal +1)
E
5 kb
Control
KDM6A
Kdm6a cKO
Control
KMT2C
Kdm6a cKO
Kmt2c
F
Kmt2c
Single cell gene expression level
Control
Kdm6a cKO
G
shKdm6a
shScr #1 #2 #3 kDa
KDM6A
H3K4me1
H3K4me2
H3K4me3
H3K27me3
Histone H3
Ponceau S.
GC1-spg
GC2-spd
Relative protein level (normalized to histone H3)
GC1
GC2
H3K4me1
H3K4me2
H3K4me3
H2K27me3
H
GSK-J4 (24 h, uM)
0 10 25 50 kDa
H3K27me3
H3K4me1
H3K4me3
H3K27ac
H3
Ex vivo testis culture
Fold Change (normalized to H3)
GSK-J4
0 μM
10 μM
25 μM
50 μM
I
GSK-J4 (24 h, uM)
rep 1 rep 2 rep 3
0 5 10 0 5 10 0 5 10 kDa
KDM6A
Beta-actin
H3K4me1
H3K4me2
H3K4me3
H3K27me3
Histone H3
GC1-spg
GC2-spd
Fold change vs. untreated (normalized to H3)
GSK-J4
0 μM
5 μM
10 μM
J
Control Kdm6a cKO
Bio. replicate 1 2 1 2 kDa
KDM6A
H3K4me1
H3K4me2
H3K4me3
H3K27me3
Histone H3
Testis
Control
Kdm6a cKO
Fold Change (normalized to H3)

**Fig. S3. Regulation of H3K4 and H3K27 methylation by KDM6A in male germ cells.**

**A,** Intersection among gene promoters with a differentially enriched KDM6A peak in *Kdm6a* cKO testis and gene promoters with a KMT2C peak detectable in control testis. **B,** RPKM-normalized signal for KMT2C ChIP-seq centered on KDM6A peaks exhibiting strong reductions (> 1.5 FC, inferred germ cell-specific) in *Kdm6a* cKO testis. **C,** ChIP-qPCR analysis for KMT2C enrichment at the *Kdm3a* gene promoter in control and *Kdm6a* cKO mixed testis. ns = non-significant by unpaired t-test. **D,** Correlation between KDM6A signal at promoter-associated peaks in testis and expression of the corresponding gene in bulk RNA-seq data from testis tissue (*26*). TPM = transcripts per million. r = Pearson correlation coefficient. **E,** Genome browser track showing ChIP-seq read alignments for KDM6A (red) and KMT2C at the *Kmt2c* locus for control and *Kdm6a* cKO testis. **F,** Gene expression for *Kmt2c* across single cells derived from testis scRNA-seq data from control and *Kdm6a* cKO mice (*26*). **G,** Western blot for the indicated histone modifications in lysates of GC1-spg (left) or GC2-spg (middle) cells expressing shScr (control) or one of three distinct sh*Kdm6a* constructs. **H,** Western blot for the indicated histone modifications in lysates of ex vivo testis cultures treated with increasing concentrations of GSK-J4, a catalytic inhibitor of KDM6A and KDM6B. **I,** Western blot for the indicated histone modifications in lysates from GC1-spg (left) and GC2-spd (right) cells treated with increasing concentrations of GSK-J4. **J,** Western blot for the indicated histone modifications and PRM2 in lysates from control and *Kdm6a* cKO caudal sperm. Ponceau Red staining total protein is shown below. For **G-J**, densitometric analysis of Western blots for the indicated histone modification normalized to total histone H3 and relative to control levels is shown at right (**G, J**) or below (**H, I**). All bars indicate mean ± SEM.

**Figure S4**

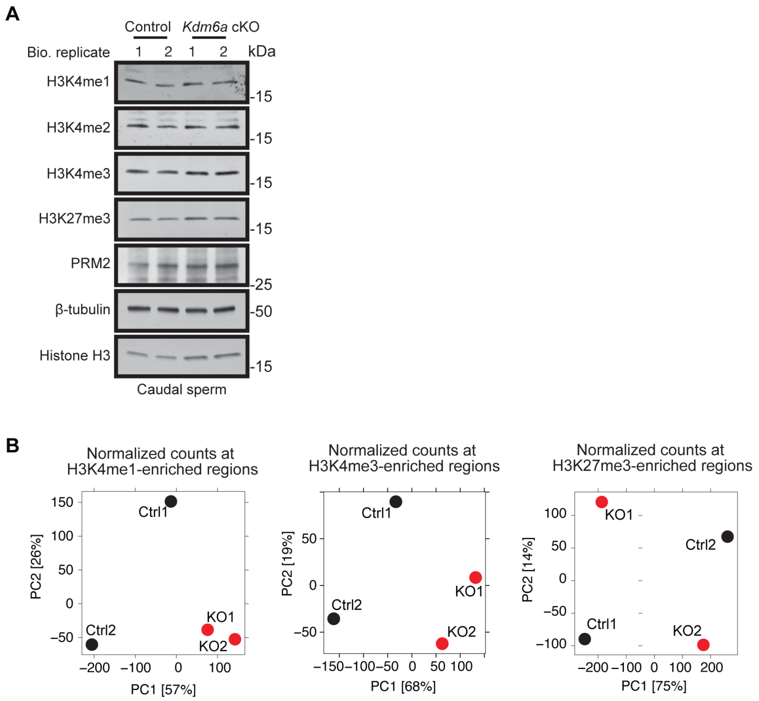


**Fig. S4. H3K4me1, H3K4me3, and H3K27me3 in *Kdm6a* cKO sperm.**

**A,** Western blot for H3K4 and H3K27 methylation, total histone H3, and protamine 2 (PRM2) in control and *Kdm6a* cKO caudal sperm. b-tubulin is a loading control. **B,** Principal component analysis of normalized ChIP-seq signal for H3K4me1 (left), H3K4me3 (center), and H3K27me3 (right) across enriched regions in sperm from control (black) and Kdm6a cKO (red) samples.

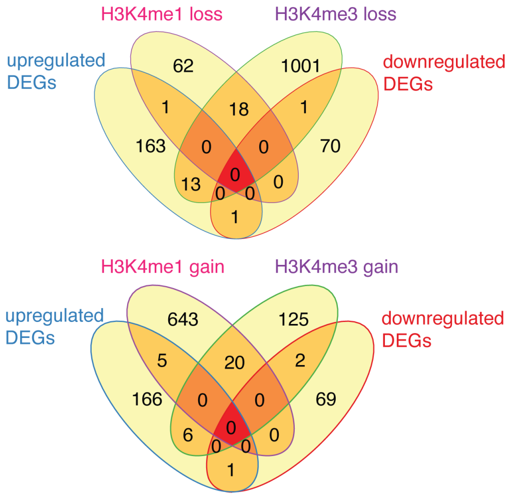


**Fig. S5. Differential gene expression overlaps in preimplantation embryos from *Kdm6a* cKO sperm.**

Overlaps of differentially expressed genes between control F1 and *Kdm6a* F1 embryos and genes associated with differential H3K4 methylation in *Kdm6a* cKO sperm relative to control.

**Figure S6**

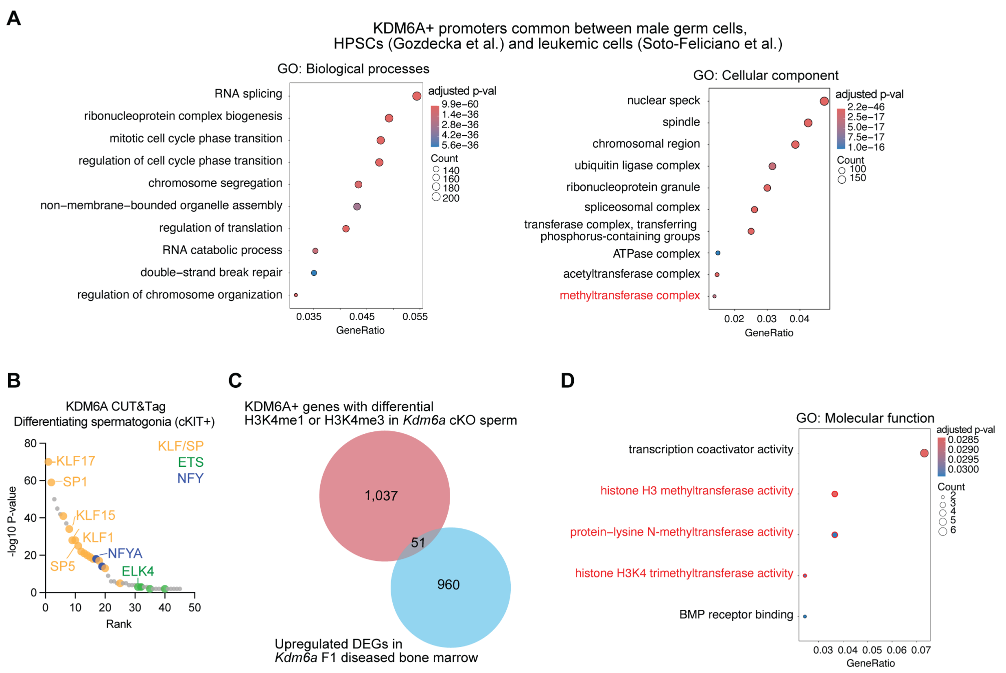


**Figure S6. Comparison of KDM6A binding and effects on histone modifications in germ and hematopoietic cells.**

**A,** Top ranked Gene Ontology terms for Biological Process (left) and Cellular Component (right) enriched among KDM6A+ promoters common across male germ cells and hematopoietic cells. **B,** DNA motifs enriched in KDM6A peaks from CUT&Tag data in sorted KIT+ differentiating spermatogonia. The three major families of transcription factors (KLF/SP, ETS, and NFY) overrepresented are highlighted and motifs common between whole testis and hematopoietic cell ChIP-seq datasets are labeled. **C,** Intersection between KDM6A+ genes with differential H3K4me1 or H3K4me3 in *Kdm6a* cKO sperm and upregulated differentially expressed genes in *Kdm6a* F1 malignant bone marrow. **D**, Enriched Molecular Function gene ontology categories among the overlapping genes from **(C).**

**Tables**

**Table S6. Antibodies used in this study**.

| Target | Manufacturer (cat #) | Application | Dilution/amount |
|---|---|---|---|
| β-Actin | Cell Signaling Technology (4970) | Western blotting | 1:5000 |
| β-Tubulin | Cell Signaling Technology (#2146) | Western blotting | 1:5000 |
| DMC1 | Proteintech (67176) | Immunofluorescence | 1:100 |
| FLAG | Sigma-Aldrich (F1804) | Western blotting | 1:3000 |
| HA | Cell Signaling Technology (3724) | Western blotting<br>Immunofluorescence | 1:3000<br>1:100 |
| H3K4me1 | Abcam (ab8895) | Western blotting<br>CUT&Tag<br>ChIP | 1:2000<br>1 µg<br>1 µg |
| H3K4me2 | Abcam (ab7766) | Western blotting | 1:2000 |
| H3K4me3 | Abcam (ab8580) | Western blotting<br>CUT&Tag<br>ChIP | 1:2000<br>1 µg<br>1 µg |
| Histone H3 | Abcam (ab1791) | Western blotting | 1:20,000 |
| KDM6A | NOVUS (NBP1-80628) | coIP | 1 µg |
| KDM6A | Cell Signaling Technology (33510) | ChIP<br>Immunofluorescence<br>Western blotting | 5 µg<br>1:100<br>1:2000 |
| KIT | R&D Systems (AF1356) | Immunofluorescence | 1:50 |
| KIT (CD117) | Invitrogen (12-1171-82) | FACS | 1 µg, 1:200 |
| KMT2C | Sigma-Aldrich (ABE1851) | coIP<br>Western blotting<br>ChIP | 3 µl<br>1:2000<br>3 µl |
| PRM2 | Briar Patch Biosciences (Hup 2B) | Western blotting | 1:2000 |
| SETD1A | Abcam , ab70378 | ChIP<br>Western blotting | 2 µg<br>1:2000 |
| VIMENTIN | Abcam, ab8978 | Immunofluorescence | 1:50 |
| Normal rabbit IgG | Sigma-Aldrich (NI01) | coIP | 3 µl |
| Donkey anti-goat Alexa Fluor 488 | Invitrogen (A11055) | Immunofluorescence | 1:500 |
| Donkey anti-mouse Alexa Fluor 488 | Invitrogen (A21202) | Immunofluorescence | 1:500 |
| Donkey anti-rabbit Alexa Fluor 568 | Invitrogen (A10042) | Immunofluorescence | 1:500 |
| Goat anti-rabbit IgG, peroxidase | Vector Laboratories (PI-1000) | Western blotting | 1:20,000 |
| Horse anti-mouse IgG, peroxidase | Vector Laboratories (PI-2000) | Western blotting | 1:20,000 |

**Table S7. Primers for qPCR**.

| Target | Forward (5’ -> 3’) | Reverse (5’ -> 3’) |
|---|---|---|
| *Hdc* | CCAGAAATAGGCCAAGGGCA | ATAGAGAAGCCAGGGCAGGA |
| *Kdm3a* | AGAAAATCGCACAGGCTCACA | GCAATCCAATAGGCGCTCA |
| *Stra8* | GGGACCTTGAAGATGGCTCC | GGCTAGCGCCAGTTCTTACA |
